# Rational engineering of lipid-binding probes via high-throughput protein-lipid interaction screening

**DOI:** 10.1101/2025.01.05.627504

**Authors:** Taki Nishimura, Kotaro Tsuboyama, Yuki Nakagaki, Eiji Yamamoto, Noboru Mizushima

## Abstract

Lipid-binding domains, originally isolated from natural proteins, are useful tools essential for analyzing membrane lipids in cells, and their applications are varied. Yet, there is no general strategy for engineering lipid-binding domains. Here, we present a robust method for monitoring protein-lipid interactions, named the Cell surface Liposome Binding (CLiB) assay. Using this technique, we isolated high-affinity lipid-binding domains and nanobodies that preferentially bind to phosphatidylinositol phosphates. Furthermore, by combining the CLiB assay with next-generation sequencing, allowing the analysis of more than 10,000 clones in parallel, we identified novel variants with enhanced binding to phosphatidylinositol phosphates and uncovered a common structural feature: a positively charged pocket necessary for binding, formed by the three loop regions. This study opens a new avenue for the rational design and generation of lipid-binding probes on demand.

## Introduction

Lipid-binding domains are protein modules, typically consisting of 45–500 amino acids, that are found in effector proteins or pathogen toxins and play a central role in the membrane association of proteins (*1*, *2*). As several lipid-binding domains specifically recognize sterols and the head groups of phospholipids and sphingolipids, they have been widely used as lipid biosensors to detect the relative abundance and subcellular localization of lipids in cells (*3*) and to evaluate the state of organelle membrane damage (*4–6*). Additionally, lipid-binding domains are employed in other applications, such as labeling apoptotic cells in flow cytometry (*7*) and isolating phosphatidylserine (PS)-positive exosomes (*8*). Thus, these domains are highly versatile and irreplaceable tools for biological research.

To date, numerous studies of lipid-binding domains have been carried out using a wide range of approaches. Structural and computational analyses have elucidated the molecular mechanisms by which lipid-binding domains interact with their target lipids and sense membrane features, including membrane curvature and charge (*1*, *9*). Systematic studies provided a broader understanding of lipid-binding domain membrane association and have partially described their lipid-binding specificities (*10–14*). Moreover, several lipid-binding domains have been engineered by introducing mutations at the membrane interface to enhance their binding affinities or to improve their binding specificities (*15–18*). However, comprehensive information on the lipid-binding affinities of lipid-binding proteins remains limited, and the fundamental principles of lipid recognition are still largely uncertain. This makes it difficult to design lipid-binding domains optimized for basic research and to predict the effects of single nucleotide polymorphisms (SNPs) within lipid-binding domains.

Most of the challenges discussed above arise from the limited throughput of current protein-lipid interaction quantification methods. In this study, we introduce a yeast display-based protein-lipid interaction assay, named the Cell surface Liposome Binding (CLiB) assay. This novel assay is compatible with high-throughput screening and next-generation sequencing analysis, expanding our capabilities for the analysis of protein-lipid interactions and the engineering of lipid-binding domains. We propose that the technique and framework presented here provide a general approach for the high-throughput quantification of protein-lipid interactions and the rational design of novel lipid-binding probes.

## Results

### Cell-surface liposome binding (CLiB) assay facilitates the analysis of protein-lipid interactions

To overcome the time-consuming and low-throughput problems in protein-lipid interaction analysis, we have developed a yeast surface display-based method, named the CLiB assay (Fig. 1A). Briefly, a lipid-binding domain was expressed on the yeast cell surface using a yeast display system. The cells were then mixed with liposomes containing the target lipids and rhodamine-conjugated phosphatidylethanolamine (PE). Protein-lipid interactions were analyzed by measuring rhodamine fluorescence of the cells using flow cytometry. The entire process, from mixing cells with liposomes to quantifying lipid-binding affinities, can be completed within 2 hours. We first checked if well-characterized lipid-binding domains retained their lipid-binding activities in the CLiB assay (*3*). Yeast cells expressing a PI3P-binding domain PX-p40phox interacted with liposomes containing PI3P in correlation with the increase of cell surface expression of PX-p40phox (Fig. 1B-D). In contrast, no binding to liposomes containing other phosphatidylinositol phosphates (PIPs) or acidic phospholipids was observed, confirming the PI3P-specific binding of PX-p40phox, as previously reported (*19–21*). Similarly, other lipid-binding domains, LactC2 and PH-PLCδ1, demonstrated preferential binding to PS and PI(4,5)P_2_, respectively (Fig. 1B-D and Fig. S1). It is known that oncogenic AKT1 mutation E17K alters the lipid-binding specificity of PH-AKT1, a lipid-binding domain for PI(3,4)P_2_ and PI(3,4,5)P_3_ (*22*, *23*). Consistent with these previous studies, the CLiB assay showed that wild-type PH-AKT1 interacted with both PI(3,4)P_2_ and PI(3,4,5)P_3_ liposomes, whereas PH-AKT1 E17K additionally bound to other PIPs species (Fig. 1B-D). These results indicate that protein-lipid interactions can be reconstituted on the yeast cell surface and that the effects of a single amino-acid point mutation on lipid-binding specificity can be assessed using the CLiB assay.

**Fig. 1.**
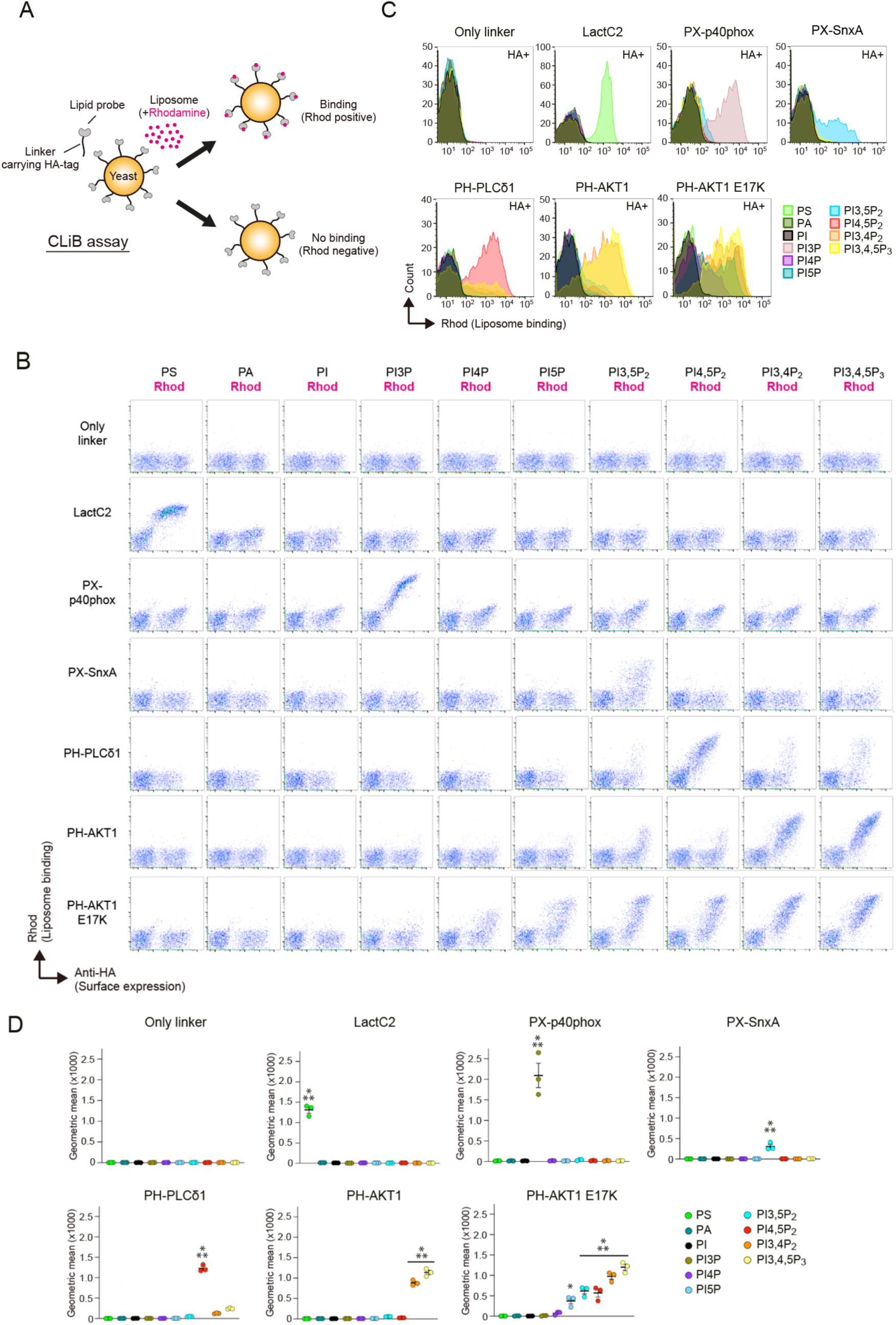
The protein-lipid interaction analysis on yeast cell surface by CLiB assay. (**A**) The cartoon showing an experimental design of CLiB assay. Yeast cells (5×10^6^ cells) expressing lipid-binding domains on their cell surfaces are incubated with the 200 μM liposomes containing the target lipids and rhodamine-conjugated PE. When the liposomes bind to the lipid-binding domains, the cells become rhodamine positive, which can be analyzed by flow cytometry. The expression levels of lipid-binding domains are assessed by anti-HA antibody staining. (**B-D**) Analysis of the lipid-binding activities of the lipid-binding domains. Wild-type SEY6210 strain was used. The indicated acidic phospholipids were contained in the liposomes as a target lipid. Two hundred μM liposomes were used as bait in CLiB assay, while 200 nM liposomes (1/1000× diluted one) were used in the case of LactC2 to minimize its non-specific binding (see Fig. S1). X axis and Y axis indicate the surface expression level and lipid-binding activity of the lipid-binding domains, respectively (B). The histogram of rhodamine signals in the HA-positive cells expressing the indicated lipid-binding domains. The threshold for HA-positive was determined using a control sample without anti-HA antibody staining (C). The geometric mean of rhodamine signals demonstrated in (C). Note that the lipid-binding domains analyzed here showed their lipid-binding specificity (D). Data represent mean ± SEM (n = 3). Differences were statistically analyzed by one-way analysis of variance (ANOVA) and Tukey multiple comparison test. Asterisks indicate statistical differences compared to samples without asterisks (\**P* < 0.05, \*\*\**P* < 0.001).

### Generation of stereospecific high-affinity clones binding to PI(3,5)P_2_ by directed evolution

To demonstrate the utility of the CLiB assay, we used it for the directed evolution of a known lipid probe. The PX domain of SnxA (PX-SnxA) was recently characterized as a PI(3,5)P_2_-binding domain with high selectivity, and its usefulness as a lipid biosensor to monitor PI(3,5)P_2_ distribution in *Dictyostelium* and mammalian cells has been demonstrated (*24*). The CLiB assay confirmed the lipid-binding ability of PX-SnxA and its specificity for PI(3,5)P_2_ (Fig. 1B-D), although its binding affinity was relatively weaker compared to other lipid-binding domains for their target lipids (geometric mean of rhodamine signals: ∼305 for PX-SnxA vs. ∼1,227 for PH-PLCδ1; Fig. 1D). To create PX-SnxA clones with high PI(3,5)P_2_-binding affinity, we prepared an error-prone PCR library of PX-SnxA with roughly three to five mutations per clone spread throughout the gene. This library was transformed into yeast cells, and the mutated PX-SnxA clones were screened for improved PI(3,5)P_2_ binders using the CLiB assay. PI(3,5)P_2_-binding clones were enriched using PI(3,5)P_2_ liposomes as bait in the first of two rounds. To exclude the less specific clones, additional liposomes containing other PIPs, such as PI3P or PI5P, were labeled with other fluorophores and used in the subsequent round, and clones showing preferential binding to PI(3,5)P_2_ liposomes were isolated (Fig. 2A). To test their functionality as PI(3,5)P_2_ biosensors in a cellular context, the isolated clones were intracellularly expressed in yeast cells, and their localization was analyzed by confocal microscopy. GFP-fused dimerized wild-type PX-SnxA did not clearly localize to the membrane structures in yeast cells, instead forming a large aggregate in the cytoplasm (Fig. 2B, upper). In contrast, seven clones isolated in the screen localized to the vacuolar membranes (Fig. 2B, lower and Fig. S2A), where PI(3,5)P_2_ is thought to be enriched (*25*). In line with this, the vacuole-localized clones exhibited higher PI(3,5)P_2_-binding activity (Fig. 2C-D and Fig. S2B) than the parental wild-type PX-SnxA clone (Fig. 1D). Clone #2179, an isolated clone, had lower cell surface expression than PX-SnxA compared to cell populations with similar levels of binding capacity (Fig. 2E), suggesting that the isolated clones were in a more functional state. To quantitatively evaluate PI(3,5)P_2_ binding affinity, the dissociation constant (Kd) value was determined based on the results of the binding quantities for different PI(3,5)P_2_ amounts in the CLiB assay (*26*) (see Methods). The vacuole-localized clones tended to have lower Kd values for the binding to PI(3,5)P_2_ liposomes, with their Kd values roughly half that of wild-type PX-SnxA (Kd 7.43 nM–16.47 nM for the isolated clones vs. 20.10 nM for wild-type; Fig. S2C). Importantly, the binding specificity for PI(3,5)P_2_ was maintained (Fig. S2B). Thus, we successfully enhanced the affinity of the PX-SnxA domain for PI(3,5)P_2_ while keeping the specificity via an unbiased, high-content CLiB screen.

**Fig. 2.**
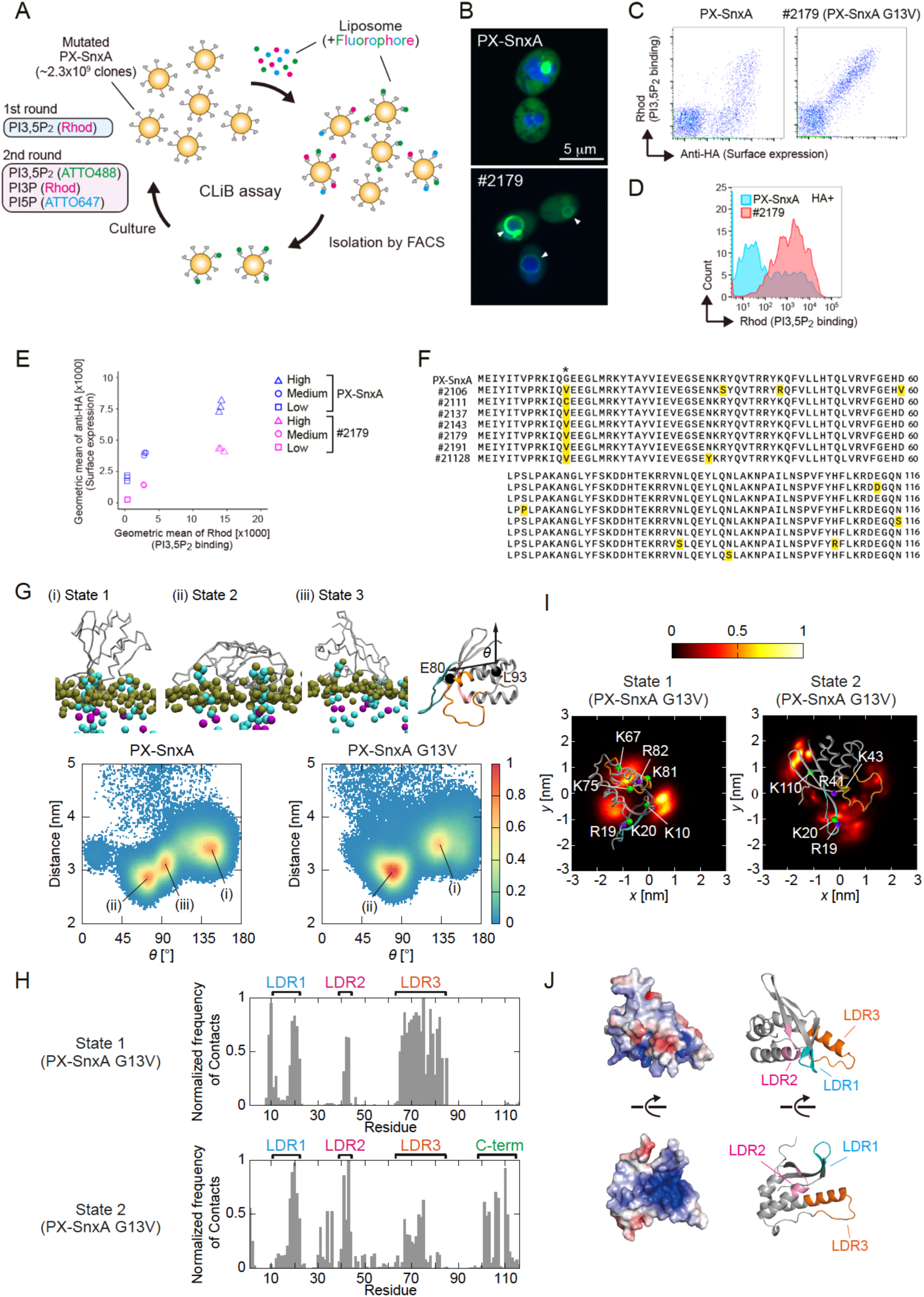
An error-prone PCR-based directed evolution of a PI(3,5)P_2_ biosensor PX-SnxA. (**A**) An experimental workflow of CLiB screen to isolate evolved clones from the mutated PX-SnxA library. In the first round, the liposomes containing PI(3,5)P_2_ and rhodamine-conjugated PE were used as bait. In the second round, three distinct liposomes were used at the same time: the first one contains PI(3,5)P_2_ and ATTO488-conjugated PE; the second one contains PI3P and rhodamine-conjugated PE; the third one contains PI5P and ATTO647-conjugated PE. The cells preferentially binding to liposomes containing PI(3,5)P_2_ were isolated by a cell sorter. (**B**) Intracellular localization of GFP-fused dimerized PX-SnxA and an isolated clone #2179 in wild-type SEY6210 cells. Vacuoles were stained with the fluorescent dye CellTracker Blue CMAC (indicated by blue). Note that the clone #2179 clearly localized to vacuolar membranes (indicated by arrowheads). Scale bar, 5 μm. (**C, D**) The PI(3,5)P_2_-binding activities of PX-SnxA and clone #2179 were measured in CLiB assay (C). Histogram of rhodamine signals of the cells expressing PX-SnxA or clone #2179 (D). (**E**) Comparison of the surface expression levels of PX-SnxA and clone #2179 showing similar lipid-binding activities. Low, Medium and High were categorized depending on the strength of rhodamine signals in CLiB assay. (**F**) Sequence alignment of the parental clone PX-SnxA and the isolated clones. Mutations introduced by error-prone PCR were highlighted by yellow boxes. The asterisk indicates the position of Gly13. (**G**) Coarse-grained molecular dynamics (CG-MD) simulations of PX-SnxA and PX-SnxA G13V interacting with the membranes. (Upper) The orientations of the PX-SnxA proteins are classified into three different representative states: state 1 (i), state 2 (ii) and state 3 (iii). Proteins are shown as silver lines, lipid phosphate groups in bronze, and PIP_2_ molecules in cyan and purple. (Lower) Normalized density map of proteins (angles relative to the membrane v.s. the *z-*component of the distance between centers of mass of the protein and the membrane). (**H**) Contact frequencies between PIP_2_ and individual amino acid residues in each state of PX-SnxA G13V. (**I**) PIP_2_ clustering in the lipid bilayer underneath bound PX-SnxA G13V. The normalized density of PIP_2_ phosphate headgroups in the lipid bilayer corresponding to each bound state is shown as a heat map. Arg (purple) and Lys (green) residues with high contact frequences (>0.7) with PIP_2_ are shown as particles. (**J**) A predicted structure of PX-SnxA by AlphaFold2 (*49*) and its representation by surface electrostatic potential. LDR1, LDR2 and LDR3 are indicated by blue, magenta and orange colors, respectively. Electrostatic surface potential map generated with the APBS plugin of PyMOL, where blue and red correspond to positive and negative electrostatic potentials, respectively.

Next, we analyzed the sequences of the evolved clones and found a high frequency of missense mutations at position Gly13 (Fig. 2F, asterisk). Of the seven clones, the glycine residue at position 13 was mutated into valine in six clones and to cysteine in one clone. One of the evolved clones, clone #2179, has the sole missense mutation, G13V, indicating that this mutation was sufficient to increase the binding affinity of PX-SnxA. To investigate how PX-SnxA and the G13V mutant interact with PI(3,5)P_2_, we performed MD simulations of the PX-SnxA proteins interacting with a PIP_2_-containing lipid bilayer and analyzed their dynamics on the membrane. Coarse-grained molecular dynamics (CG-MD) simulations followed our previously validated protocol shown to effectively identify PIP-binding modes for various membrane-bound proteins (*27*). The structures of PX-SnxA and PX-SnxA G13V were predicted using AlphaFold. The CG-MD simulations of 10×10 μs were conducted for each system, with the protein initially positioned away from the membrane in various orientations. The protein diffused in the aqueous environment and quickly bound to the membrane, adopting three distinct binding modes with varying orientations and distances relative to the membrane (Fig. 2G). In one of the bound states, the loop regions — named lipid-binding determining regions LDR1, LDR2, and LDR3 (Fig. 2H) — were oriented toward and bound to the membranes (state 1; Fig. 2G-H and Fig. S2D). In the other states, the proteins adopted a lying-down configuration on the membranes (states 2 and 3; Fig. 2G-H and Fig. S2D). The wild-type PX-SnxA exhibited reversible dynamics, maintaining equal distribution among states 1, 2 and 3 (Fig. S2E). In contrast, the G13V clone irreversibly transitioned from the upright position (state 1) to a lying-down state (state 2) (Fig. S2E), forming additional interactions with the membranes via its C-terminus (Fig. 2H). These findings suggest that state 2, which predominates in the G13V clone, represents a high-affinity PI(3,5)P_2_ binding state of the PX-SnxA proteins.

To better understand the molecular interactions between PX-SnxA G13V with PIP_2_, we analyzed the density of the PIP_2_ headgroup within the membrane for each bound state, as well as its interactions with individual amino acid residues. Our analysis revealed that the phosphate groups of PIP_2_ are positioned in close proximity to specific basic residues (Arg19, Lys20, Arg40, Arg41, Lys43, Lys67, and Arg82) within the LDRs (Fig. 2H-I). Moreover, the surface electrostatic potentials demonstrated that the three LDRs formed a positively charged pocket (Fig. 2J). Together, these results suggest that the three LDRs serve as a positively charged binding pocket critical for PI(3,5)P_2_ recognition, and that the G13V mutation modifies the conformational preferences of PX-SnxA configuration on membrane surfaces.

### Deep mutational scanning analysis of lipid-binding domains

By combining the CLiB assay with next-generation sequencing (NGS), we dramatically increased the throughput of protein-lipid interaction analysis. To demonstrate the capacity of the high-throughput CLiB (HT-CLiB) assay, we conducted a deep mutational scanning (DMS) analysis for PX-SnxA G13V, PX-p40phox, PH-PLCδ1, and PH-AKT1, which comprehensively investigate how individual residues contribute to the lipid-binding ability of these four lipid-binding domains. First, we synthesized an oligo pool containing lipid-binding domains with all possible single amino acid substitutions. After generating the yeast display library (Fig. 3A, i), the cells were incubated with liposomes containing the target lipids (Fig 3A, ii). Subsequently, four fractions were collected using a cell sorter according to their rhodamine signals (Fig. 3A, iii). To assess the quality of cell sorting, we reanalyzed the sorted fractions using the CLiB assay and confirmed the different lipid-binding strengths of each fraction as expected (Fig. S3A and S3B). We then evaluated the lipid-binding abilities of individual clones by counting each clone in all fractions using NGS analysis (Fig. 3A, iv-v). Two replicate results from the SnxA G13V DMS library showed a strong correlation (R^2^ = 0.98) (Fig. 3B), confirming the high reproducibility of the HT-CLiB assay.

**Fig. 3.**
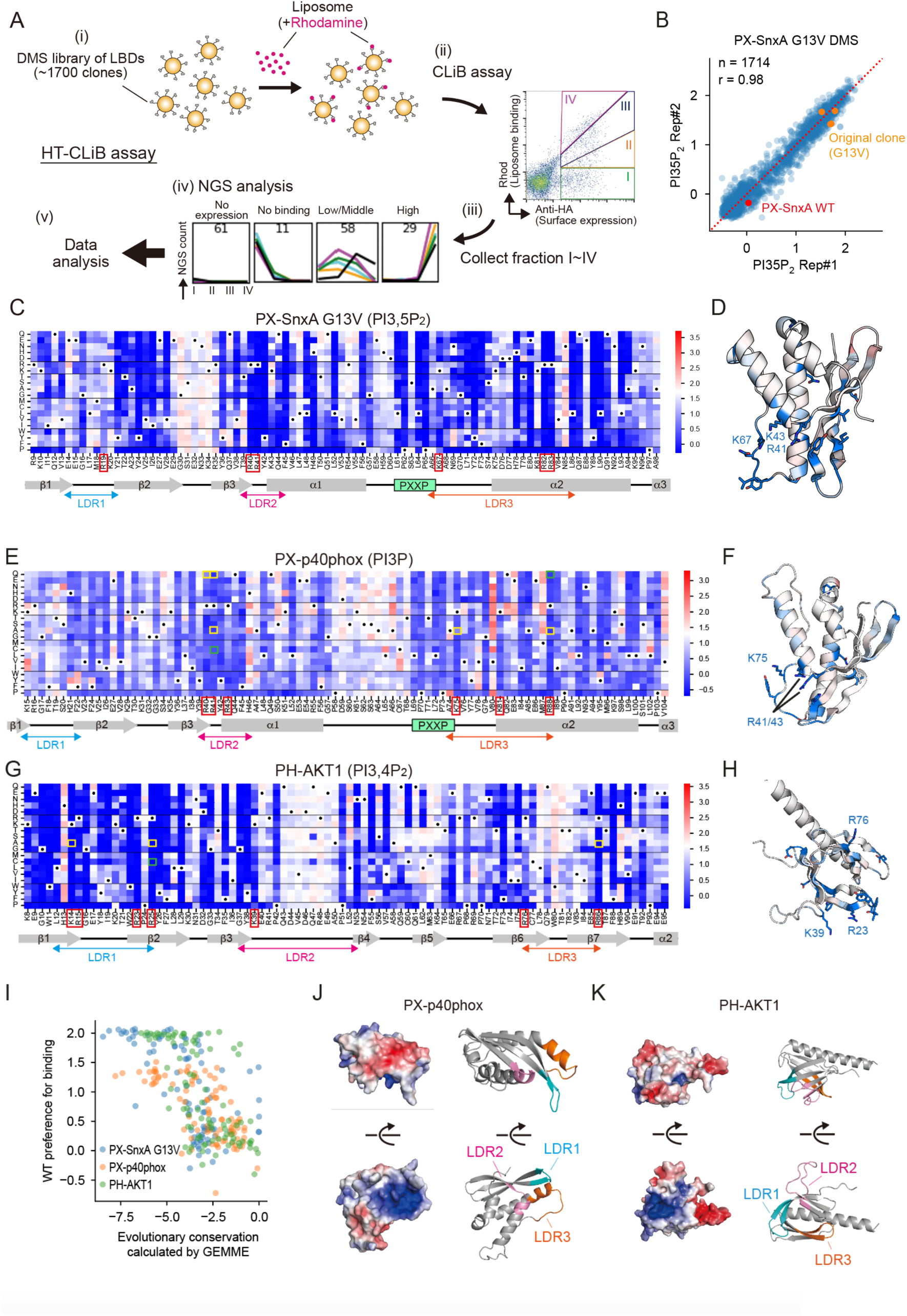
Comprehensive mutation analysis of lipid-binding domains reveals a key feature for their PIPs binding activity. (**A**) The series of experiments to carry out high-throughput CLiB (HT-CLiB) assay. The deep mutation scanning (DMS) library of the lipid-binding domains was designed and synthesized, and then inserted into a plasmid for yeast display system (i) (*34*). The DMS library was mixed with the liposomes containing the target lipids and rhodamine-conjugated lipids (ii, CLiB assay). After staining with anti-HA antibody, the cells were sorted by a cell sorter according to their lipid-binding activities (iii). The plasmids were purified using a yeast plasmid miniprep kit from the cell pellets, followed by PCR to amplify the coding regions of lipid-binding domains for the NGS analysis. NGS count was calculated and demonstrated as line graphs of the individual clones. Each color indicates a lipid binding activity against the individual lipids (iv). Relative lipid-binding activities of the individual clones based on their NGS counts are calculated and demonstrated as a heatmap (v, shown in Fig. 4C-H). (**B**) A scatter plot showing the correlation between the HT-CLiB results derived from two independent analyses of PI(3,5)P_2_ binding activities of the PX-SnxA G13V DMS library. (**C-H**) The HT-CLiB results for the libraries of PX-SnxA G13V (C, D), PX-p40phox (E, F), and PH-AKT1 (G, H). Heat maps show the binding activity scores of each mutant (see Methods section). White indicates wild-type binding activity, and red and blue indicate strong and weak binding activities, respectively. Black dots indicate wild-type amino acids (C, E and G). Their predicted structures are colored by the median of binding activity score changes at each position. Only the surface residues (burial <1.5; see Methods section for the detail) were colored. AlphaFold2 model (*49*) was used for their structural predictions (D, F and H). (**I**) A plot showing the relationship between wild-type preference for the lipid-binding and the evolutionary conservation of each amino acid. (**J, K**) A predicted structure of PX-p40phox (J) and PH-AKT1 (K) and their surface electrostatic potential. LDR1, LDR2 and LDR3 are indicated by blue, magenta and orange colors, respectively. Electrostatic surface potential map generated with the APBS plugin of PyMOL, where blue and red correspond to positive and negative electrostatic potentials, respectively.

The DMS analysis successfully quantified the affinity score (see Methods section for the score calculation), which for each mutant is shown in heatmaps (Fig. 3C-H and Fig. S3C-D). Consistent with previous reports (Table S1), various known lipid-binding deficient mutants showed a significant reduction in lipid-binding abilities in the DMS analysis (Fig. 3E, 3G, and Fig. S3C, highlighted by yellow boxes). In the AKT1 DMS library, the binding affinity for PI(3,4)P_2_ was strongly correlated with that for PI(3,4,5)P_3_ (Fig. S3E), indicating that most AKT1 clones recognize the phosphate groups at the 3rd and 4th positions on the inositol ring. Next, we individually analyzed notable mutants that drastically altered lipid-binding affinity in the HT-CLiB assay and found that these clones showed either lower or higher affinities for their target lipids than the original clones, as shown in the DMS results (Fig. S3F-I). This series of results indicate that our large-scale analysis is relatively quantitative and useful for isolating clones of desired binding strengths.

We also found a positive correlation between the preference for wild-type (WT) amino acids and amino acid sequence conservation among species, as calculated by GEMME (*28*) (Fig. 3I), suggesting that key residues involved in lipid binding (i.e., function of the domains) are more likely to be conserved during evolution. Additionally, the binding-affinity changes observed in the HT-CLiB assay are closely related to the impact of disease-related single nucleotide polymorphisms (SNPs) in previous studies. For example, chronic granulomatous disease-related mutations in the *NCF4* gene encoding the p40phox (R41C and R88Q) (*29*, *30*) showed less binding activity to PI3P (Fig. 3E, highlighted by green boxes). *AKT1* R25C, which has been identified as a pathogenic mutation related to Cowden syndrome (*31*, *32*), reduced the PI(3,4)P_2_-binding affinity of AKT1 (Fig. 3G, highlighted by a green box). Thus, our DMS dataset could serve as an important reference for inferring the effects of SNPs on the lipid-binding ability of lipid-binding domains.

DMS analysis, in agreement with the MD simulation results (Fig. 2G-I), further revealed that Lys and Arg residues located in LDR1-3 are required for full lipid-binding activity of PX-SnxA G13V (Fig. 3C, red boxes), although many residues distributed in the central part of the structure also contributed (Fig. 3C-D). Consistent with this, the corresponding regions in PX-p40phox were critical for the PI3P-binding activity of PX-p40phox (Fig. 3E, red boxes and Fig. 3F), indicating that the three LDR regions are commonly required for PIPs binding. To determine the general mechanisms beyond the PX domain, we examined PH domains that possess distinct backbone structures. The interstrand loops (indicated as LDR1-3 in Fig. 3G and Fig. S3C) vary significantly in sequence and structure between the PH domains in different species, although they are commonly involved in target lipid recognition (*27*, *33*). The DMS analysis revealed that Lys and Arg residues in the LDRs are essential for the PIPs-binding activity of PH-AKT1 (Fig. 3G, red boxes) and PH-PLCδ1 (Fig. S3C, red boxes, and S3H) as well as the PX domain shown above. In line with this, these PIPs-binding domains contained a positively charged pocket enclosed by the three LDRs (Fig. 3J-K, and S3J). Most Lys/Arg residues were found at both ends (N/C terminals) of each LDR, reflecting that they were located at the core of a pocket to add positive charges. Collectively, we concluded that the positively charged pocket formed by the three LDRs is a key feature commonly found in PIPs-binding domains.

### Isolation and characterization of lipid-binding nanobodies

Similar to the three LDRs of PIPs-binding domain involved in recognition of the target lipids, nanobodies, heavy-chain antibodies derived from camelids, also bind to target molecules via three complementarity-determining regions (CDRs) (Fig. 4A). Based on this analogy, we used the CLiB assay to isolate PIPs-binding nanobodies from a synthetic nanobody library. This library consisted of approximately 1×10^8^ unique clones (*34*). After several rounds of the CLiB assay screening, we obtained various clones that bound to liposomes containing PIP_2_ species, such as PI(3,5)P_2_, PI(3,4)P_2_ or PI(4,5)P_2_. Some of these isolated clones promiscuously interacted with all PIP_2_ and PS lipid species (Fig. S4A), while others showed preferential binding to few PIP_2_ species (Fig. S4A, see clones #7, #26 and #28).

**Fig. 4.**
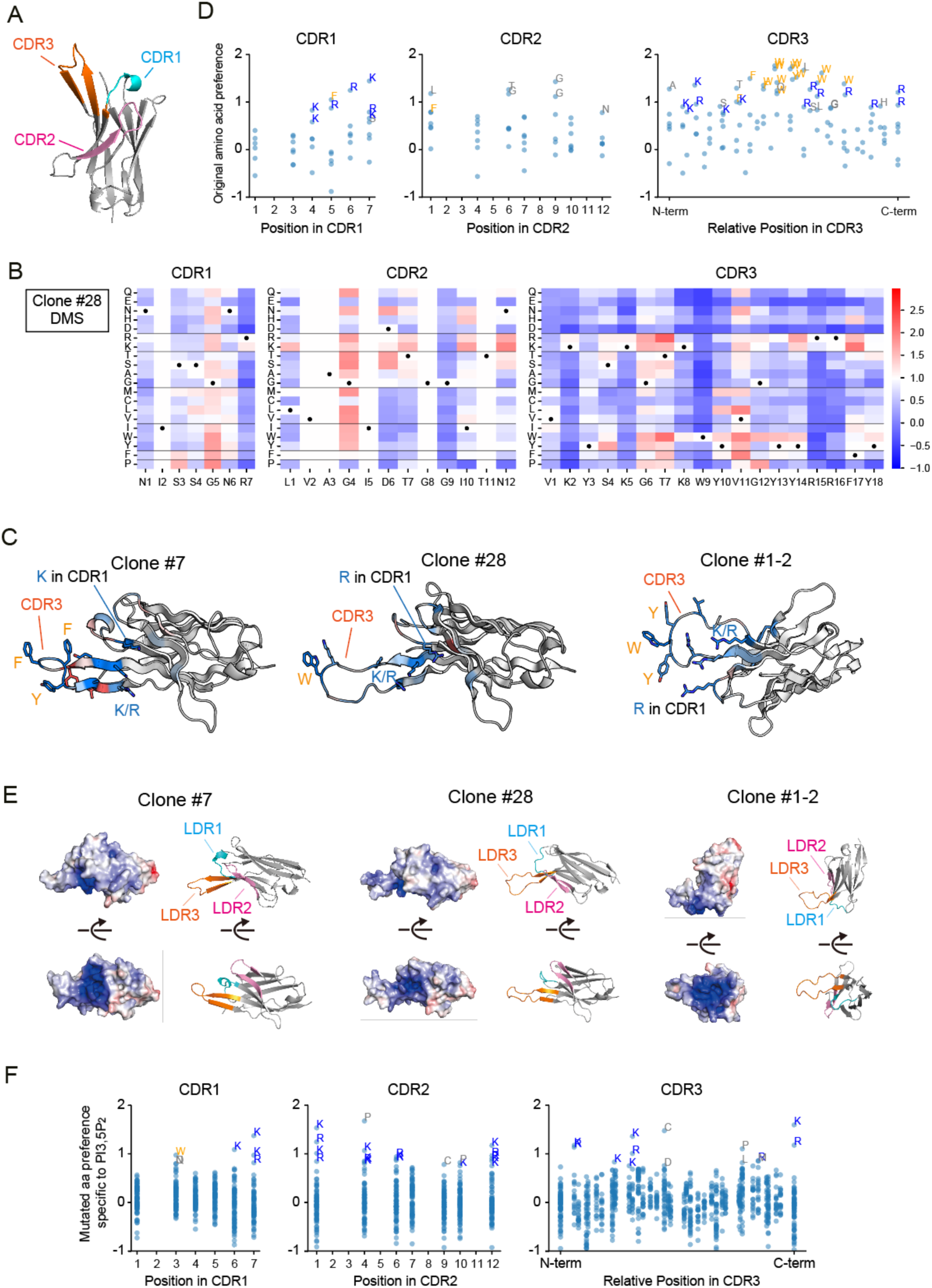
Isolation and analysis of PIPs-binding nanobodies. (**A**) An example of a predicted nanobody structure by AlphaFold2. (**B**) Mutational scanning results for Nb clone #28. Heat maps show the binding activity scores of each mutant. White indicates wild-type binding activity, and red and blue indicate strong and weak binding activities, respectively. Black dots indicate wild-type amino acids. Note that we did not replace the amino acids at position 2 in CDR1 and positions 2, 3, and 8 in CDR2 because they are fixed in the original nanobody library (*34*). (**C**) The predicted structures of clones #7, #28 and #1-2, colored by the median of binding activity score changes at each position. AlphaFold2 model (*49*) was used for their structural predictions. **(D)** Preference of original amino acids of the isolated nanobodies for each position. Note that basic amino acids at C-terminal in CDR1 and at N-/C-terminal in CDR3, and bulky aromatic amino acids in the middle of CDR3 are strongly preferred. (**E**) The predicted structures of the indicated nanobody clones and their surface electrostatic potential. LDR1, LDR2 and LDR3 are indicated by blue, magenta and orange colors, respectively. Electrostatic surface potential map generated with the APBS plugin of PyMOL, where blue and red correspond to positive and negative electrostatic potentials, respectively. (**F**) Preference of mutated amino acids for each position. Replacement to basic amino acids at C-terminal in CDR1 and N-/C-terminal in CDR2 and CDR3 increase the binding affinity to PI(3,5)P_2_.

To investigate the causal relationship between amino acid residues and the preferential binding to all PIP_2_ species, we conducted DMS analysis of CDR1-3 regions for the 22 isolated nanobody clones in a single HT-CLiB assay. The total number of sequences analyzed was approximately 12,000. After checking the quality of cell sorting (Fig. S4B and S4C), each fraction was subjected to NGS analysis, and the lipid-binding affinities of each clone were calculated (Fig. 4B and 4C) as described above (Fig. 3A). The DMS analysis of seven nanobodies (showing (i) the original sequence affinity was not saturated, and (ii) the DMS pattern did not show strong bias among PIP_2_ species) and the original amino acid preferences (Original clone binding score - median of 19 mutants at the position) were also calculated (Fig. 4D). In CDR1, a preference for basic amino acids was observed in the C-terminal region. This tendency was also observed in the N- and C-terminal regions of CDR3. Additionally, the central region of CDR3 strongly preferred bulky hydrophobic residues, such as Trp and Phe (Fig. 4C and 4D, see CDR3). Since the C-terminus of CDR1 and both termini of CDR3 are in close proximity (Fig. 4A and 4C), we speculated that the polarized distribution might reflect a structural requirement for PIPs-binding. Indeed, according to their predicted structures, all isolated PIPs-binding nanobodies contained a positively charged pocket that is flanked by CDRs (Fig. 4E). Using DMS data of four nanobodies (Nb #6-11, Nb #1-28, Nb #2-201A, and Nb #3A-30; Fig. S4A) showing mutations that cause differential preferences between PS and PIP_2_ species, we further investigated what mutations change the binding preferences for PI(3,5)P_2_ and PS, and found that basic residues in the C-terminus of CDR1 and both sides of CDR2 and CDR3 increase preferential binding to PI(3,5)P_2_ (Fig. 4F and Fig. S4D). These results suggest that the binding preference of nanobodies for PIP_2_ is influenced by the positive charge in the C-terminal region of CDR1, and the N-/C-terminal regions of CDR2 and CDR3.

### *In silico* design and improvement of nanobodies via a machine learning-based approach

We tested whether a large dataset from DMS of PIPs-binding nanobody clones could be applied to a machine learning approach to achieve *in silico* design. A prediction model for the lipid-binding activity of the nanobodies was constructed. Briefly, the amino acid sequence of each clone was transformed into feature values using the protein language models, UniRep (*35*) or ESM-2 (*36*), and the eXtreme-Gradient-Boosting (XGBoost)-based top model was trained using 90% of the DMS data from the 22 nanobody clones measured by the CLiB assay (Fig. 5A, left). The lipid-binding prediction model was validated using test data that were not used for the model training. The performances of the UniRep-based and ESM-2-based prediction models were almost comparable (r = 0.87 vs 0.86; Fig. 5B). In the second step, we performed Markov chain Monte Carlo-based *in silico* directed evolution using the prediction model (*37*) (Fig. 5A, right): essentially, (i) a single mutation was randomly introduced into the CDR1-3 of ∼10,000 parental sequences, which were included in the synthetic nanobody yeast display library (*34*), (ii) then the mutational effect predicted via the prediction model, (iii) and finally mutations were selected for the next cycle where, neutral or beneficial mutations were always accepted while sometimes slightly detrimental mutations accepted so that the algorithm was allowed to explore a larger amino acid sequence space (see Methods for details). This mutation and prediction cycles were repeated 500 times.

**Fig. 5.**
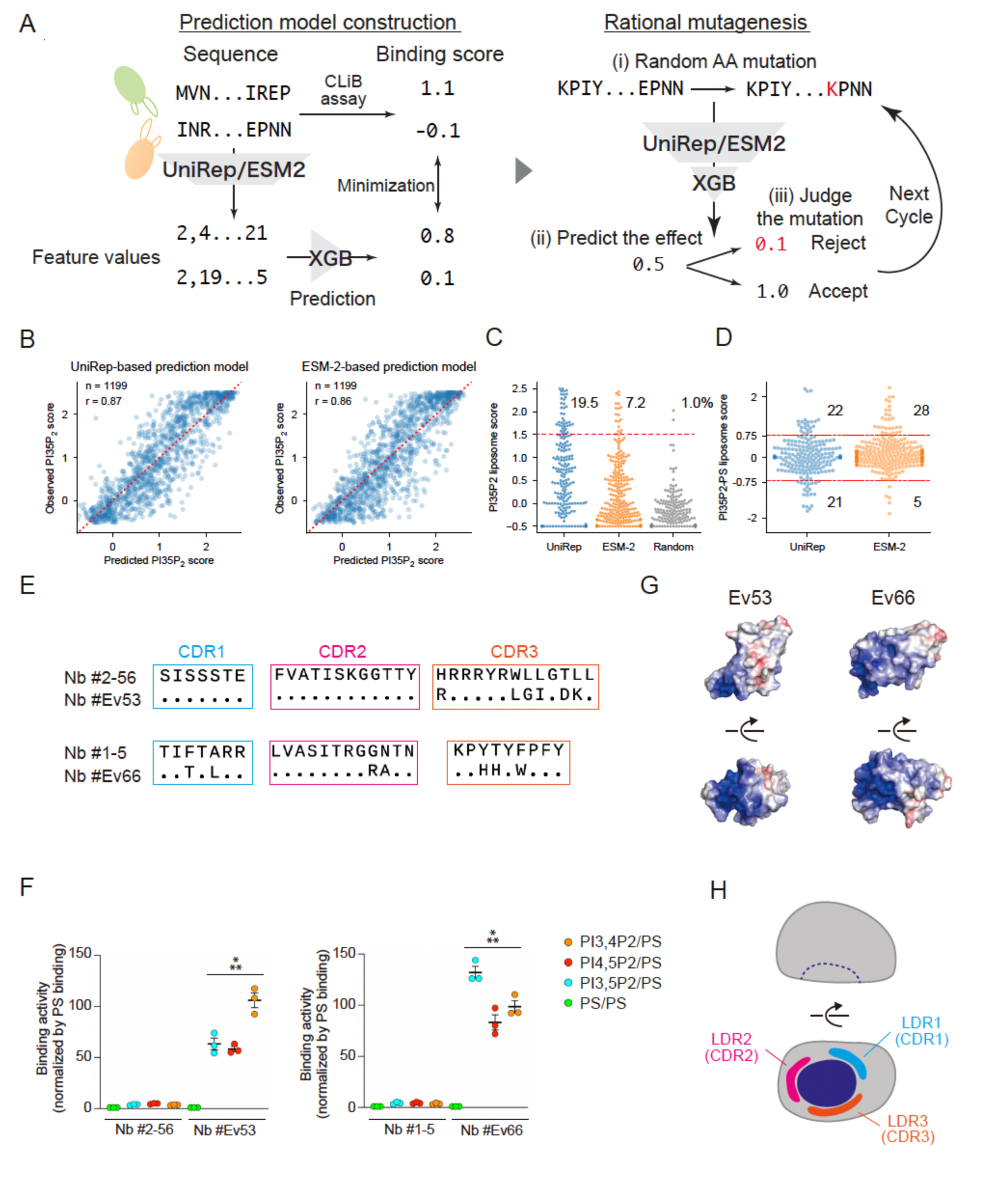
*In silico* design and directed evolution of PIPs-binding nanobodies. (**A**) The workflows for prediction model construction (left) and rational mutagenesis (right). (**B**) Performance of UniRep-based (left) and ESM-2-based (right) prediction models on binding affinity against PI(3,5)P_2_. (**C**) Binding affinity score distribution of nanobodies designed by UniRep-based, ESM-2-based prediction model and randomly mutated nanobodies. Percentage of the clones showing high binding scores for each group is shown on the top. The red dashed line represents binding score = 1.5 and the percentiles left to the beeswarm plot represent the fraction of nanobodies with score above the threshold. (**D**) Binding specificity score (PI(3,5)P_2_ [target liposome] binding score - PS [non-target liposome] binding score) distribution of nanobodies designed by UniRep-based and ESM-2-based design. The numbers right to beeswarm plots represent the number of nanobodies with expected (PI(3,5)P_2_-oriented) and unexpected (PS-oriented) specificity (binding specificity score > 0.75 or < -0.75, which are represented by red dashed lines). (**E**) Sequence alignment of *in silico* evolved clones (Nb #Ev53 and Nb #Ev66) and their parental clones (Nb #2-56 and Nb #1-5), respectively. (**F**) The lipid-binding preferences of the indicated clones. Lipid-binding activity was analyzed in CLiB assay. The BJ5465 protease-deficient strain was used. To investigate PIPs preferences, the geometric mean of rhodamine signals against the indicated lipids were normalized by that against PS. As for Nb #2-56 and Nb #1-5, data shown in Fig. S4A were reanalyzed. Data represent mean ± SEM (n = 3). (**G**) The surface electrostatic potentials of evolved nanobody clones based on their predicted structures using AlphaFold2 model. Electrostatic surface potential map generated with the APBS plugin of PyMOL, where blue and red correspond to positive and negative electrostatic potentials, respectively. (**H**) A cartoon showing a concept of PIPs-binding pocket that is organized by basic residues located in the bottom regions of three loops. LDR1(CDR1), LDR2(CDR2) and LDR3(CDR3) are indicated by blue, magenta and orange colors, respectively. A dark blue area indicates a positively charged pocket. Differences were statistically analyzed by one-way analysis of variance (ANOVA) and Tukey multiple comparison test. Asterisks indicate statistical differences compared to samples without asterisks (\*\*\**P* < 0.001).

To rationally design PI(3,5)P_2_-binding nanobodies, we tested two protein language models, UniRep and ESM-2. From ∼10,000 amino acid sequences generated by this rational mutagenesis approach, we filtered 200-250 nanobody clones that are predicted to exhibit strong binding affinity to PI(3,5)P_2_. After validating the purity of the sorted cells (Fig. S5A, B), the lipid-binding strength of each clone was analyzed using NGS. Compared to random amino acid replacement, both UniRep-based and ESM-2-based mutagenesis were more likely to generate nanobodies with a strong affinity for PI(3,5)P_2_ compared to random mutagenesis (19.5% [UniRep], 7.2% [ESM-2] vs. 1.0% [random]; Fig. 5C). ESM-2-based designs exhibited greater specificity for PI(3,5)P_2_ liposomes (vs. PS liposomes) (Fig. 5D), although UniRep-based mutagenesis was more likely to generate PI(3,5)P_2_ binders than ESM-2-based mutagenesis (Fig. 5C).

Although the error-prone PCR-based directed evolution of nanobodies did not efficiently improve their lipid-binding specificity (data not shown), we found that two clones generated by ESM-2-based design demonstrated their preferential binding to PIP_2_ species including PI(3,5)P_2_ rather than PS compared to the parental clones (Fig. 5E-F). This indicates the advantage of our *in silico* design approach over error-prone based directed evolution (Fig. 2). Additionally, consistent with the nanobody clones that were isolated in the CLiB screen, these clones contained an additional arginine residue in CDR2 or CDR3 (Fig. 5E) and a positively charged pocket formed by CDRs (Fig. 5G). This suggests that ESM-2-based mutagenesis also ‘understand’ that the formation of a concave structure with a positive charge is a prerequisite and likely sufficient for PIPs-binding, regardless of the protein backbone structures (Fig. 5H).

## Discussion

In this study, we demonstrated that protein-lipid interactions can be analyzed on the yeast cell surface using the CLiB assay (Fig. 1). As the yeast display system is inherently compatible with directed evolution and high-throughput analysis, the CLiB assay has innovatively extended our capabilities for lipid-binding domain engineering and enabled several breakthroughs: (i) the isolation of PX-SnxA clones through directed evolution that bind efficiently to PI(3,5)P_2_ with high affinity and specificity (Fig. 2), (ii) comprehensive quantification of affinities for individual lipid species from over 10,000 single mutants of natural lipid-binding domains (Fig. 3) and nanobodies (Fig. 4) through the HT-CLiB assay, and (iii) *in silico* design and directed evolution of PIP_2_-specific nanobodies (Fig. 5). To the best of our knowledge, this study is the first to present lipid-binding data for more than 10,000 different proteins and predict lipid-binding activity based on amino acid sequence information.

We found that PIPs-binding proteins have one thing in common: a positively charged pocket. Given that the head group of PIPs is relatively larger than that of other negatively charged lipids, such as PS and phosphatidic acid, proteins require sufficient space at the lipid-binding site to accommodate PIPs without steric hindrance. Our data indicate that having three loops (LDRs and CDRs) is geometrically optimal for forming a pocket that retains diversity. The sizes and net charges of the pockets will depend on the amino acid composition and secondary structures of the loops. In addition, we showed that high-affinity nanobody clones possess bulky hydrophobic residues at the center of CDR3, suggesting that the hydrophobic wedge formed by these residues facilitates its interaction with membranes, followed by the recognition of the PIP lipid head group of by a pocket via the electrostatic interactions. However, the lipid-binding specificities of the isolated nanobody clones against individual PIP species were suboptimal compared to those of the lipid-binding domains analyzed in this study, implying the existence of additional key factors influencing binding specificity. These differences should be addressed in future studies.

This method, however, has some limitations. Lipid-binding domains carrying many cysteine residues, such as the FYVE domain and PKC-C1 domains, did not show specific binding to their target lipids in the CLiB assay, contrary to previous reports (*3*), likely due to their structural instability in our yeast display system. The protein size for the HT-CLiB assay is also constrained by the maximum length of the DNA oligo pool synthesis. Additionally, it is important to consider that large-scale measurements and automated data processing have the potential to introduce inaccuracies, although we have confirmed that some key results (such as mutations that enhance lipid-binding affinities) were reproducible in the single clone experiments (Fig. S3).

We envision that the CLiB assay will serve as a basic platform for compiling comprehensive datasets, including lipid-binding affinities for millions of lipid-binding domains and protein scaffolds (including nanobodies), while also advancing the understanding of the fundamental principles of protein-lipid interactions. Importantly, this strategy could be applied to the development of lipid-binding probes for other lipid species. Given that the lack of reliable lipid biosensors for lipids has long been a long-standing issue in cell biology, this technology and its products are expected to see widespread use in the near future. Thus, the CLiB assay will present a promising avenue for the rational design of lipid-binding probes.

## Acknowledgements

We thank K. Karasawa and K. Kiyokawa for technical assistance with wet experiments. We thank Hiroyuki Noji, Naohiro Terasaka, Takahiro Kosugi, Toyoshi Fujimoto, Toshiki Itoh, Christopher J. Stefan, Andy Lam and members of Nishimura lab and Tsuboyama lab for the helpful discussion. We would like to thank Editage for English language editing.

## Author contributions

T.N. conceived the project and developed the CLiB assay. T.N. and K.T. designed the research. T.N. and K.T. performed most experimental work and analyzed the data. T.N. performed wet experiments and the imaging analysis. K.T. performed NGS analysis, oligo pool design and computational studies. Y.N. and E.Y. designed, performed, and analyzed the MD simulation. T.N., K.T., E.Y., and N.M. acquired funding. T.N. and K.T. wrote the original draft and revised the manuscript with input from all authors.

## Data and materials availability

All data and codes for reproducing the figures will be released upon the official publication.

## Methods

### Plasmids

Descriptions of plasmids and primers used in this study are in Table S3. pYDS649 vector was isolated from a clone derived from Yeast-Display Nanobody Library (EF0014-FP, Kerafast) (*34*), and its NheI restriction site was replaced with SalI by the conventional cloning method. DNA sequences encoding LactC2 (aa 270-427, NP_788783.1), PX-p40phox (aa 19-140, NP_000622.2), PX-SnxA (aa 61-175, XP_636026.1), PH-PLCδ1 (aa 11-140, NP_058731.2), PH-AKT1 (aa 1-120, NP_001014431.1), and PH-AKT1 E17K were synthesized by Twist Bioscience. They were amplified by PCR and subcloned into the BamHI/SalI site of the pYDS649 plasmid. The directed evolved PX-SnxA clones were subcloned into pRS426 plasmid containing GMAP-210 coiled-coil region (aa 39-377, NP_004230.2) and EGFP (*38*). All the point mutations in the PX-SnxA G13V, PX-p40phox, PH-AKT1, and PH-PLCδ1 were introduced by a conventional PCR-based method.

### Liposome preparation

Materials used in this study are listed in Table S3. Lipids were mixed in the desired molar ratio and the organic solvent was dried under nitrogen gas and vacuumed through a rotary evaporator (MV-100, TOMY) for 20 min at room temperature. The lipid film was rehydrated and resuspended in a selection buffer containing 20 mM Hepes-NaOH pH 7.5, 150 mM NaCl, 0.1% BSA, and 5 mM Maltose, with vortexing for 30 sec. To make liposomes (50∼60 nm diameter), the suspensions were sonicated on ice using a tip sonicator (Q500 sonicator) for 10 sec ON/10 sec OFF cycles for a total of 10 min at 25% amplitude concurrently. Lipid debris was removed by 17,700×*g* for 20 min at 4°C. The liposomes were kept at 4°C and were used within a few days.

### CLiB assay

Yeast cell lines used in this study are listed in Table S4. Standard techniques were used for yeast growth and manipulations. SEY6210α or BJ5465 strains were cultured at 30 °C for a day in YND-Trp medium (0.17% yeast nitrogen base without amino acids and ammonium sulfate, 0.5% ammonium sulfate, and 2% glucose) supplemented with synthetic drop-out medium without tryptophan (US Biological, D9531). Two hundred μL of the cell suspension was added to 2 mL of YNGal-Trp medium containing 2% galactose instead of 2% glucose to induce surface display. After the culture for 24 h, 0.5OD (∼5×10^6^ cells) cells were taken into a low-binding tube (WATSON, PK-15C-500N), washed with 300 μL of the selection buffer (20 mM Hepes-NaOH pH 7.5, 150 mM NaCl, 0.1% BSA, and 5 mM Maltose) and centrifuged at 3,000×*g* for 1 min at 4°C. After aspiration of the supernatant, the cell pellets were mixed with 40 μL of 0.5 mM unlabeled liposomes containing 70 mol% of phosphatidylcholine (PC) and 30 mol% of phosphatidylethanolamine (PE) for blocking and incubated for 30 min at 25°C with continuous shaking in a Thermo BIOSAN TS-100C shaker at 800 rpm. Twenty μL of 0.2 mM liposomes containing 68 mol% of PC, 29.5 mol% of PE, 2 mol% of the target lipids and 0.5 mol% rhodamine-conjugated PE were added, and incubated for a further 15 min on the shaker. The samples were centrifuged at 3,000×*g* for 1 min at 4°C, and the supernatant was aspirated. The cell pellets were mixed with 10 μL of Alexa 647-conjugated anti-HA antibody (MBL M180-A64, 1/200× diluted with the selection buffer) and incubated on ice for 15 min. After centrifugation and removal of the supernatant, the cell pellets were resuspended with 500 μL of the selection buffer and analyzed by flow cytometry with a FACSymphonyA1 cell analyzer (BD Bioscience). The acquired data was analyzed using FlowJo software version 10.10.0. The dissociation constant Kd values were calculated by nonlinear fit assuming one site - specific binding in GraphPad Prism 10 software.

### Generation of mutated PX-SnxA library by error-prone PCR

To prepare an insert, the PX-SnxA gene was mutagenized with error-prone PCR using a JBS Error-Prone Kit (Jena Bioscience, PP-102) in a total volume of 10 μL. Four μL of the PCR product was amplified using a Quick Taq HS DyeMix (TOYOBO, DTM-101) in a total volume of 200 μL, purified using a gel purification kit (Nippon Genetics, FG-91302), and eluted with 30 μL water. To prepare a vector, six μg of pYDS649 plasmid was digested by BamHI/SalI enzymes, purified and eluted with 30 μL water. Yeast electroporation was conducted according to the protocol reported by Benatuil et al. (*39*). In brief, BJ5465 strain was cultured at 30 °C for a day in 5 mL YPD medium (1% yeast extract, 2% bacto peptone and 2% glucose). An aliquot of the culture was inoculated into 100 mL of YPD media to give an OD600 of ∼0.3. The cells were grown until OD600 reached ∼1.6 before collecting by centrifugation. The cell pellets were washed twice with cold water, once with the electroporation buffer (1 M sorbitol/1 mM CaCl_2_), resuspended with 0.1 M lithium acetate/10 mM dithiothreitol, and incubated at 30 °C for 30 min with shaking. After centrifugation at 3,000×*g*, the cell pellets were washed with the electroporation buffer, then suspended in 100∼200 μL of the electroporation buffer to reach 1 mL volume. This corresponds to ∼1.6×10^9^ cells/mL. Three hundred fifty μL of the cell suspension was electroporated with the insert and vector DNAs by 2.5 kV using an electroporator (NEPA porator, NEPA Gene). After the electroporation, cells were suspended in 10 mL of 1:1 mix of 1 M sorbitol: YPD media and incubated at 30 °C for 1 h with shaking. The cells were collected, resuspended with 10 mL of YND-Trp medium, and spread on 33 YND-Trp plates using 300 μl of cell suspension per plate. The number of transformants was determined by plating 10-fold serial dilutions of transformed cells on YND-Trp plates and counting colonies after 3 days. Add 1 mL of water on the plates, collect the colonies with a scraper and repeat the process again. After centrifugation, the cell pellets were resuspended with YND-Trp medium to give a concentration of ∼2×10^9^ cells/mL. Finally, the cell suspension was mixed with 1/10× volume of dimethyl sulfoxide (DMSO) and stored at -80°C until analysis.

### Generation of yeast display libraries using oligo pools

Oligo pools containing all single possible mutants of PX-SnxA G13V (1,730 clones), PX-p40phox (1,710 clones), PH-PLCδ1 (1,710 clones), PH-AKT1 (1,672 clones) and nanobodies (11,986 clones) were synthesized by Twist Bioscience. As for *in silico* designed clones, an oligo pool containing nanobodies (2,100 clones) was synthesized. The oligo pools were solubilized with water to make a stock solution (∼10 ng/μl). Two μL of the 1/10× diluted oligo pools (1 ng/μL) were amplified using KOD-one (TOYOBO, KMM-101) in a total volume of 25 μL. To add the 5’ and 3’ regions and increase the amount, forty μL of 1/10× diluted 1st PCR products were amplified using KOD-one in a total volume of 1 mL. Subsequently, the PCR products were purified using a gel purification kit and eluted with 30 μL water. The suitable number of PCR cycles was chosen based on the results analyzed by MultiNA (Shimadzu) to avoid over-amplification. Yeast electroporation and library storage were conducted as described above.

### Isolation of PIPs-binding clones by CLiB assay

Fifty μL (∼0.9×10^8^ cells) of yeast display library stock was taken, inoculated in 10 ml of YND-Trp medium, and cultured at 30 °C for a day. One mL of the cell culture was added to 10 mL of YNGal-Trp medium. After the culture for 24 h, 10OD (∼1×10^8^ cells) cells were taken into a low-binding tube (WATSON, PK-15C-500N), washed with 1 mL of the selection buffer and centrifuged at 3,000×*g* for 1 min at 4°C. After aspiration of the supernatant, the cell pellets were mixed with 400 μL of 0.5 mM unlabeled liposomes containing 70 mol% of PC and 30 mol% of PE for blocking and incubated for 30 min at 25°C with continuous shaking in a Thermo BIOSAN TS-100C shaker at 800 rpm. Two hundred μL of 0.2 mM liposomes containing 68 mol% of PC, 29.5 mol% of PE, 2 mol% of the target lipids and 0.5 mol% rhodamine-conjugated PE were added, and incubated for a further 15 min on the shaker. The samples were centrifuged at 3,000×*g* for 1 min at 4°C, and the supernatant was aspirated. The cell pellets were mixed with 40 μL of Alexa 647-conjugated anti-HA antibody (MBL M180-A64, 1/200× diluted with the selection buffer) and incubated on ice for 15 min. After centrifugation and removal of the supernatant, the cell pellets were resuspended with 8 mL of the selection buffer. Rhodamine and Alexa 647 labeled cells were sorted using a cell sorter (Sony, SH800S) equipped with a 100 μm sorting chip and collected in a 15 mL tube containing 2 mL of YND-Trp medium. The cytometer was set to Ultra Purity mode. For high-throughput CLiB assay, four fractions were collected by a cell sorter according to their rhodamine signals as no-, low-, medium- and high-binding fractions (Fig. 3A). To ensure a sufficient data size, cells corresponding to more than 400 times the original diversity were sorted. In the case of the CLiB assay using multiple liposomes, no staining with anti-HA antibodies has been carried out. After cell sorting, the cells were centrifuged, resuspended with 2 mL of YND-Trp medium and cultured at 30°C for 2 days. Nine hundred μL of cell suspension was mixed with 100 μL of DMSO and stored at -80°C until analysis. The remaining cell suspension (∼1 mL) was used for next-generation sequencing (NGS) analysis.

### DNA oligonucleotide library construction

All DNA sequences were reverse-translated and codon-optimized using DNAworks2.0 (*40*). Most sequences were optimized using E. coli codon frequencies although we utilized yeast cells. All libraries were purchased from Twist Bioscience.

### Next-generation sequencing sample preparation

For NGS library preparation, the extracted plasmid from yeast cells was amplified by PCR using KOD one (TOYOBO) to add P5 and P7 NGS adapters and sequencing sequences. The number of PCR cycles was chosen based on a test qPCR run to avoid over-amplification. The DNA fragment length was confirmed in Agarose gel or MultiNA (Shimadzu), then samples were analyzed by the Miseq or Novaseq 6000 system (Illumina).

### Processing of next-generation sequencing data

Each NGS library was de-multiplexed by a unique 8-nt barcode sequence. The paired reads from each NGS library were combined by the PEAR program (*41*), then the adapter sequences were removed by Cutadapt (*42*). Reads were considered counts for a sequence only if the read perfectly matched the ordered sequences at the nucleotide level.

### Calculating binding scores from next-generation sequencing data

The fraction of each sequence was calculated from the raw counts for each sample (No, Low, Medium, High fraction, see Fig3A). Then binding score of each sequence was calculated according to the formula X:

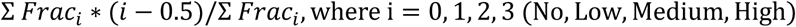

### Prediction model of lipid-binding affinities

We constructed a prediction model as in the previous studies (*35*, *37*, *43*). We utilized two protein language models, UniRep and ESM-2, to generate embeddings from amino acid sequences of nanobodies. In UniRep-based embedding, the “h_avg” was generated with the default parameters/weights. In ESM-2-based embedding, mean embeddings for the final layer from “esm2_t36_3B_UR50D” model were used.

As our top layer model to predict lipid-binding affinities, we utilized an eXtreme Gradient Boosting (XGBoost) model. We used the default parameters of python xboost library, with the exception that the maximum depth of layers is limited to three to prevent over-fittings.

### *In silico* directed evolution of lipid-binding nanobodies

We basically followed the protocol shown in the previous study (*37*, *43*). Essentially, we tried to use an algorithm that can find more functional (i.e. stronger affinities to a desired lipid type) on average but not forced to do so to find global maximum but not local maximum. To this end, we applied a Metropolis–Hastings Markov chain Monte Carlo algorithm.

Our *in silico* directed evolution was performed as follows:

1, Prepare The initial sequence(s) of nanobodies and a top model that predicts binding affinities from UniRep or ESM-2 embeddings as the inputs for the pipeline.

2, Predict binding affinity scores (y_0) of initial sequences using the top model.

3, Introduce one amino acid mutation to each of the initial sequences in CDR1-3 except position 2 in CDR1 and positions 2, 5, 6, 8, and 11 in CDR2 and predict binding affinity scores (y_1) of the mutated amino acid sequences.

4, Calculate acceptance probability according to the following formula:

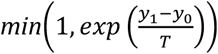

where T is temperature.

5, Judge if the mutation is accepted or not according to the acceptance probability. 6, Iterate steps 3-5 for a predetermined number of iterations.

Essentially, in the *in silico* directed evolution method, we always introduce mutations that can keep or improve the binding affinities, and we also introduce a part of mutations that weakly reduce affinities to explore larger amino acid sequence space. To this end, in the prospective evolution of lipid-binding nanobodies, ∼11,000 evolutionary trajectories were run in parallel for 1,000 iterations. We set temperature at 0.1 for the first 900 iterations, 0.01 for 900-950 iterations, 0.001 for the last 50 iterations to explore a large space of amino acid sequences but still evolve amino acid sequences whose affinities were improved.

Then we filtered 200 (for UniRep-based and random mutagenesis) or 250 nanobodies (for ESM2-based mutagenesis) for experiments from ∼11,000 sequences after 1,000 iterations according to the affinity scores predicted by the top model.

### MD simulation

Coarse-grained molecular dynamics (CG-MD) simulations were performed using GROMACS 2022.6 with the Martini 2.2 force field (*14*, *44*). The symmetric lipid bilayer used in the simulations was constructed via CHARMM-GUI (*45*), containing 340 DOPC (69%), 150 DOPE (30%), and 6 PIP_2_ (1%) molecules. Each leaflet of the bilayer was composed of 170 DOPC, 75 DOPE, and 3 PIP_2_ molecules. The PIP_2_ was modified from the Martini lipid POP2 by replacing C2B with D2B to model the fatty acid tails as 18:1.

The clone #2179 was built using Modeller, with the PX domain of SnxA as a template predicted by AlphaFold. The proteins were placed 15 nm above the lipid bilayer, and ∼25,500 water molecules along with 150 mM NaCl ions were added to the system. Simulations were conducted 10 times for each protein under different initial orientations. All systems underwent 1,000 steps of energy minimization followed by a 1 ns *NPT* constant simulation for equilibration. Ten independent production runs of 10 μs each were performed, using a timestep of 20 fs. An elastic network was applied with a cutoff range of 0.5–0.9 nm. All simulations were performed at a temperature of 310 K and a pressure of 1 bar. The Parrinello-Rahman semi-isotropic barostat was used for pressure control (*46*), and velocity rescaling was employed for temperature control (*47*).

For the contact frequency analysis, a contact was defined when the distance was below 0.7 nm.

### Amino acid burial calculation

Burial values and contact counts were computed based on AlphaFold models. The burial calculation was done as in the previous report (*48*). In brief, to calculate the burial or contacts of residue X, we added up the number of residues in a cone projecting out 9 Å away from the Cβ atom on residue X in the direction of the residue X Cα-Cβ vector. ‘Burial’ indicates the number of Cα atoms in the cone.

### Fluorescence Microscopy

The yeast cells were stained with 0.1 mM CMAC dye to visualize the vacuole. Images were obtained with a 60× PlanAPO oil-immersion lens (1.42 NA; Olympus), using a confocal FV3000 confocal laser microscope system (Olympus). All images were collected as square images with 2048 × 2048 pixels. For the final output, images were processed using ImageJ (v1.54f) in Fiji and Adobe Photoshop 2025 v26.0.0 software (Adobe).

### Statistical analysis

Differences were statistically analyzed by one-way analysis of variance and Tukey multiple comparison test. Statistical analysis was carried out using GraphPad Prism 10 (GraphPad Software). All data are presented as the means ± SEM. Reproducibility of all results reported here was confirmed.

### Use of AI tools

We utilized ChatGPT, DeepL, DeepL Write and Claude to improve the English text and the coding.

## Abbreviations

PI(3,5)P_2_: phosphatidylinositol 3,5-bisphosphate
PIPs: phosphatidylinositol phosphates
PE: phosphatidylethanolamine
PS: phosphatidylserine
MD: molecular dynamics
PX domain: phox domain
PH domain: pleckstrin homology domain
Nb: nanobody
CDR: complementarity-determining region
DMS: deep mutational scanning

---

**Fig. S1.**
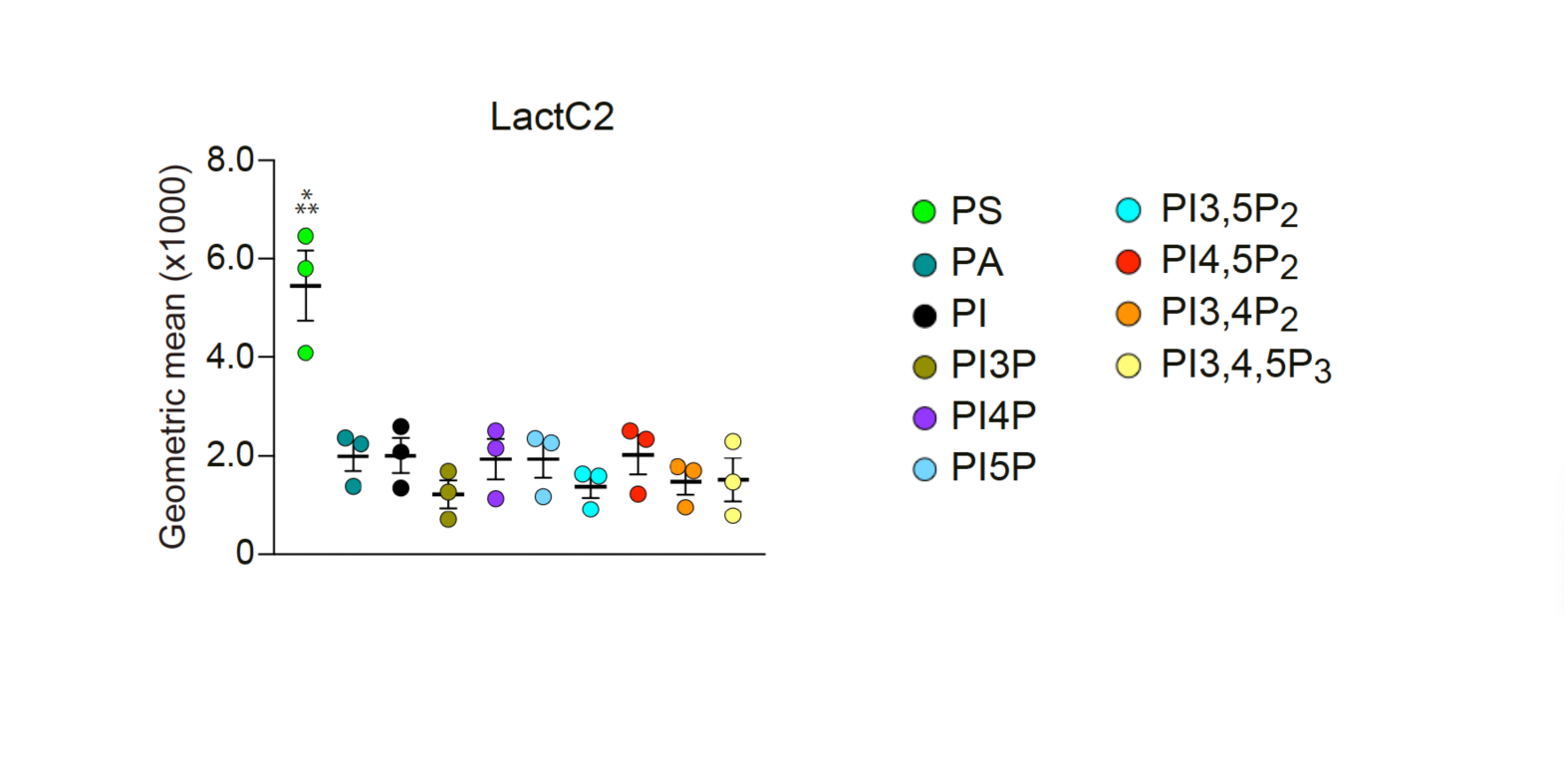
The lipid-binding analysis of LactC2 by CLiB assay. Wild-type SEY6210 strain cells expressing LactC2 were analyzed in CLiB assay. Two hundred μM liposomes containing the indicated lipid species were used as bait. The geometric means of rhodamine signals of HA-positive cells were calculated. Note that LactC2 preferentially interacted with PS compared to the other lipid species. Data represent mean ± SEM (n = 3). Differences were statistically analyzed by one-way analysis of variance (ANOVA) and Tukey multiple comparison test. Asterisks indicate statistical differences compared to samples without asterisks (\*\*\**P* < 0.001).

**Fig. S2.**
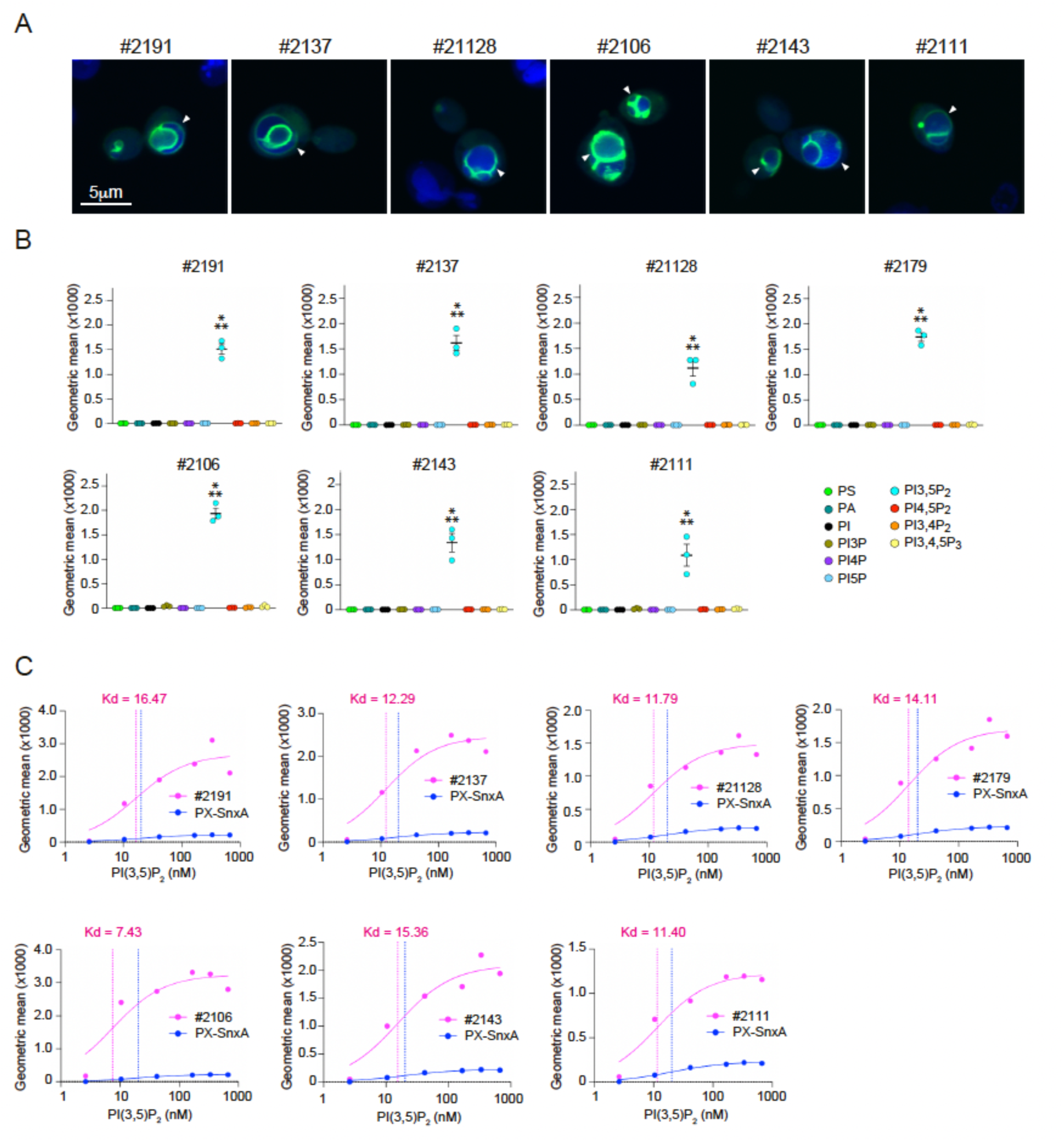
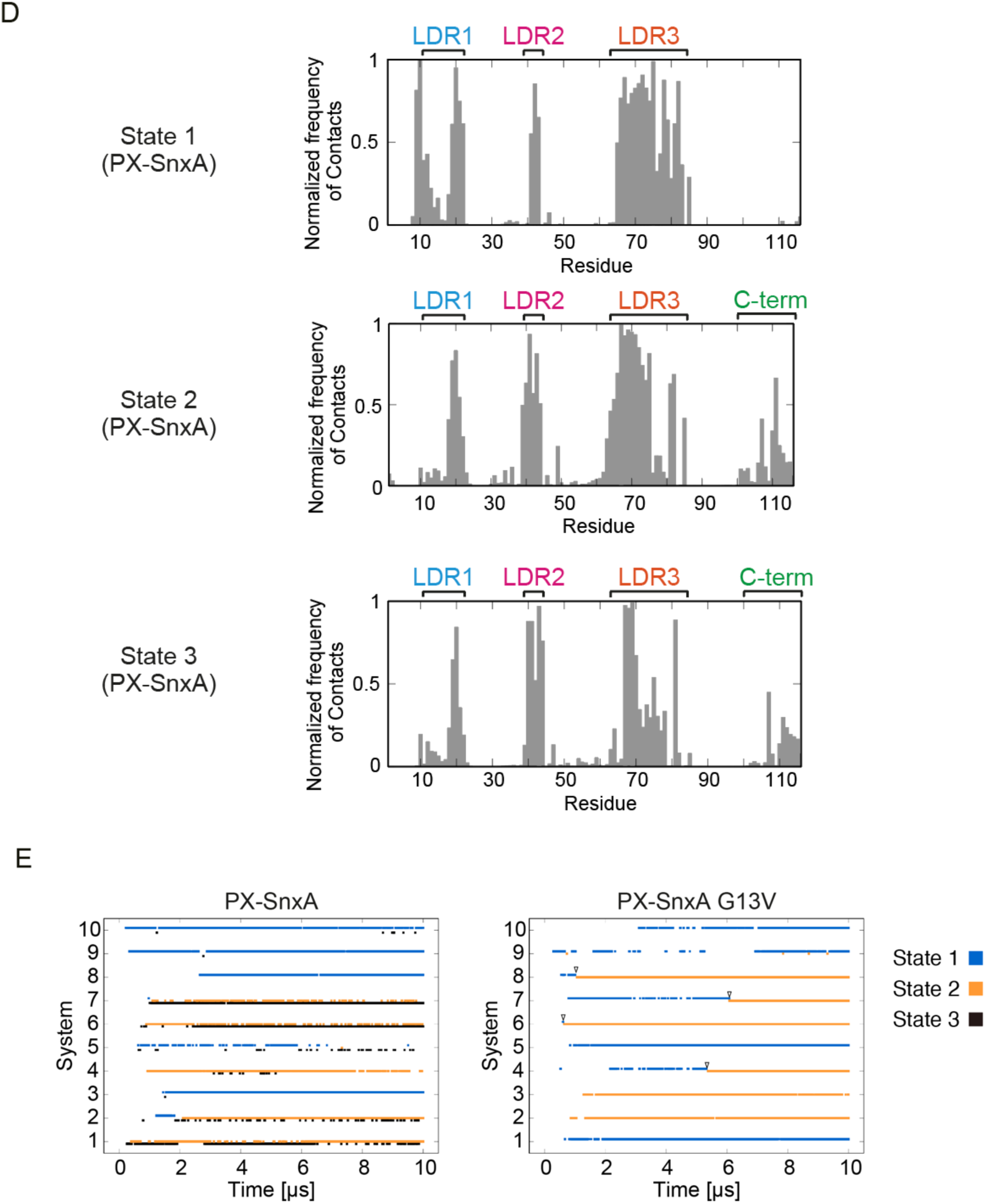
Analysis of PX-SnxA evolved clones. (**A**) Intracellular localization of GFP-fused dimerized the isolated clones in wild-type SEY6210 cells. Vacuole was stained with the fluorescent dye CellTracker Blue CMAC (indicated by blue). Note that the isolated clones localize to vacuolar membranes (indicated by arrowheads). Scale bar, 5 μm. (**B**) The analysis of lipid-binding activities of the isolated clones by CLiB assay. Data represent mean ± SEM (n = 3). Note that all clones showed their PI(3,5)P_2_-specific binding activities in CLiB assay. (**C**) PI(3,5)P_2_ binding activity against various PI(3,5)P_2_ concentrations analyzed by CLiB assay. The Kd values were determined from their binding curves (see Methods). (**D**) Contact frequencies between PIP_2_ and individual amino acid residues in each state of PX-SnxA. (**E**) Temporal variation in the membrane-binding states of PX-SnxA and PX-SnxA G13V during CG-MD simulations. State 1, state 2 and state 3 are indicated by blue, orange and black, respectively. Arrowheads indicate the time positions of the state changes of PX-SnxA G13V from state 1 to state 2. Differences were statistically analyzed by one-way analysis of variance (ANOVA) and Tukey multiple comparison test. Asterisks indicate statistical differences compared to samples without asterisks (\*\*\**P* < 0.001).

**Fig. S3.**
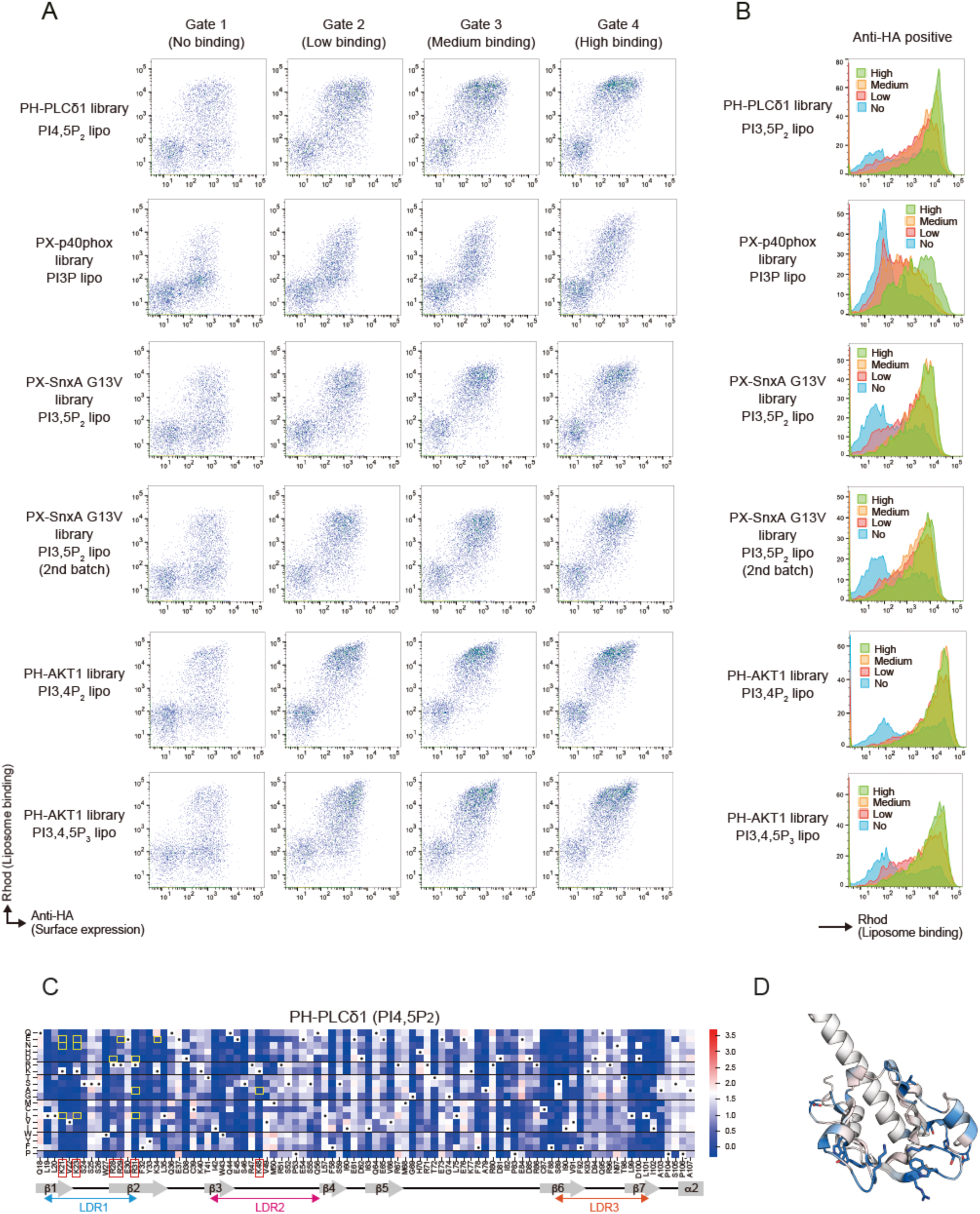
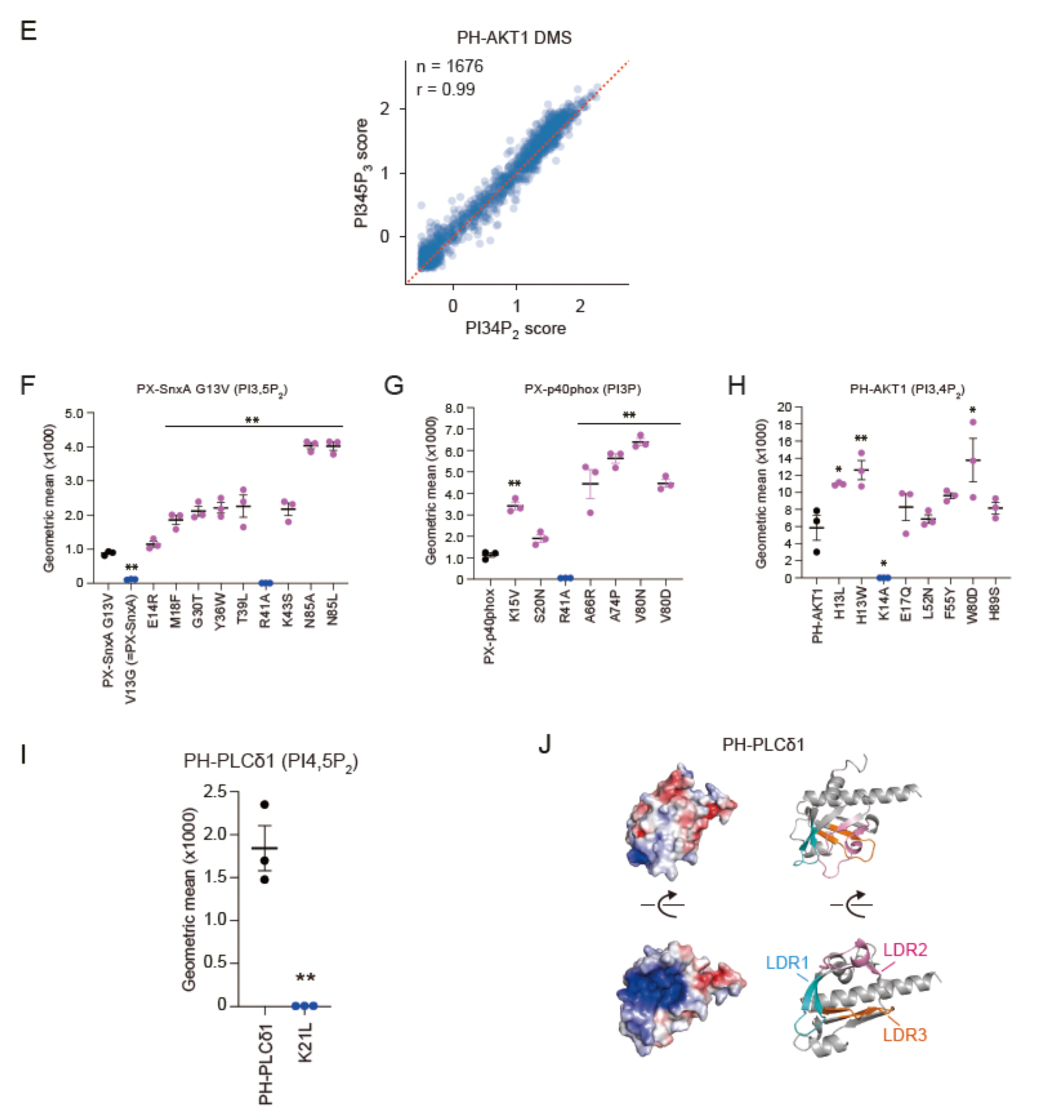
Comprehensive mutation analysis of lipid-binding domains reveals the contribution of individual amino acid residues for their lipid-binding activities. (**A, B**) Analysis of the lipid-binding activities of the sorted fractions in CLiB assay using the indicated DMS libraries and liposomes. X axis and Y axis indicate the surface expression level and lipid-binding activity of the lipid-binding domains, respectively (A). The histogram of rhodamine signals in the HA-positive cells expressing the indicated lipid-binding domains (B). (**C**) The HT-CLiB result for the libraries of PH-PLCδ1. A heat map shows the binding activity scores of each mutant. White indicates wild-type binding activity, and red and blue indicate strong and weak binding activities, respectively. Black dots indicate wild-type amino acids. (**D**) The predicted structure of PH-PLCδ1 colored by the median of binding activity score changes at each position. (**E**) A scatter plot showing the correlation between the HT-CLiB results of PI(3,4)P_2_ and PI(3,4,5)P_3_-binding activities of the PH-AKT1 DMS library. (**F-I**) The lipid-binding analysis of the mutants of PX-SnxA G13V (F), PX-p40phox (G), PH-AKT1 (H), and PH-PLCδ1 (I) by CLiB assay. The BJ5465 protease-deficient strain was used. Blue and red indicate clones low- and high-binding affinity clones shown in Fig. 3 and Fig. S3C, respectively. Data represent mean ± SEM (n = 3). (**J**) A predicted structure of PH-PLCδ1 and its surface electrostatic potential. LDR1, LDR2 and LDR3 are indicated by blue, magenta and orange colors, respectively. Electrostatic surface potential map generated with the APBS plugin of PyMOL, where blue and red correspond to positive and negative electrostatic potentials, respectively. Differences were statistically analyzed by one-way analysis of variance (ANOVA) and Tukey multiple comparison test. Asterisks indicate statistical differences compared to samples without asterisks (\**P* < 0.05, \*\**P* < 0.01).

**Fig. S4.**
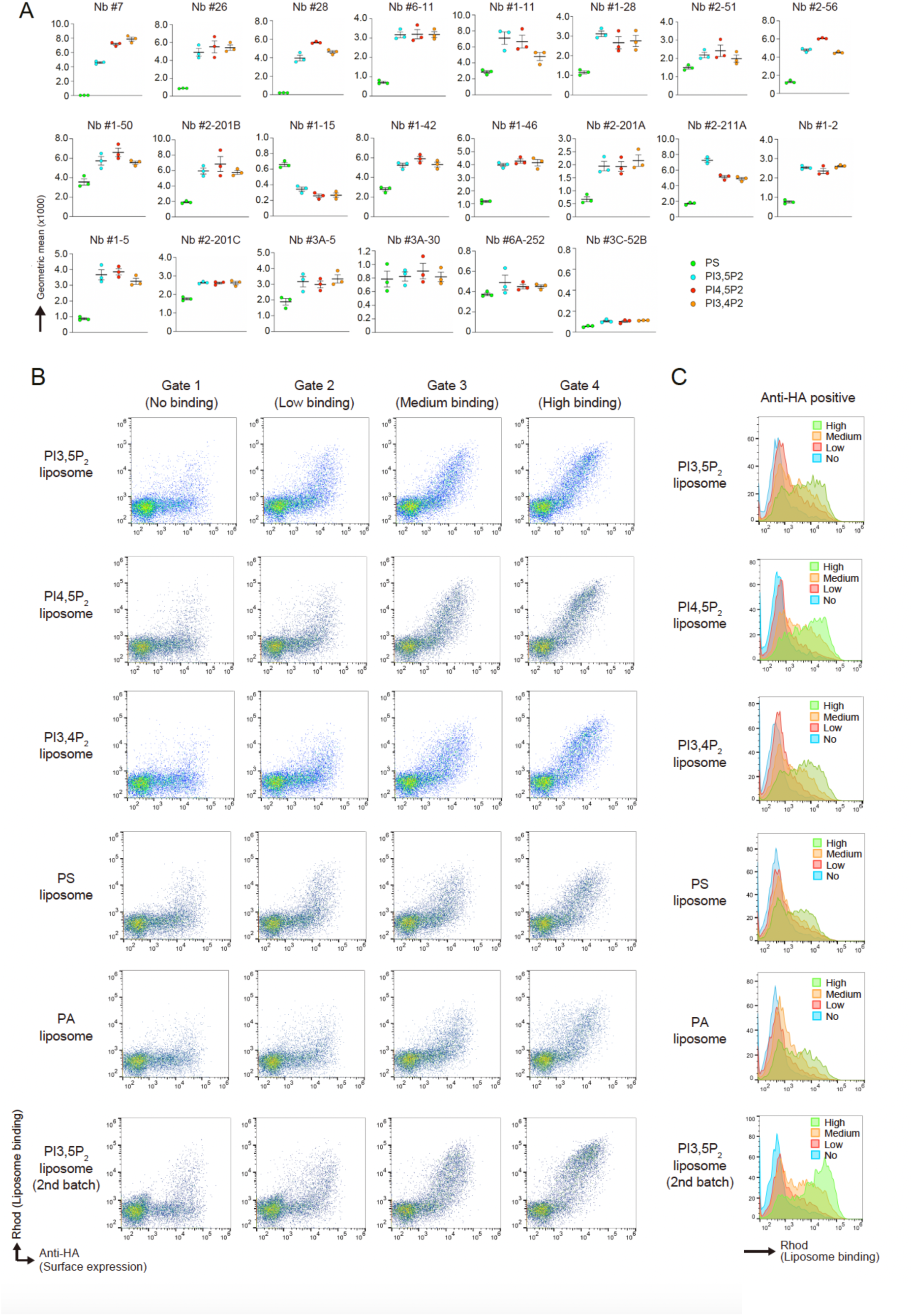
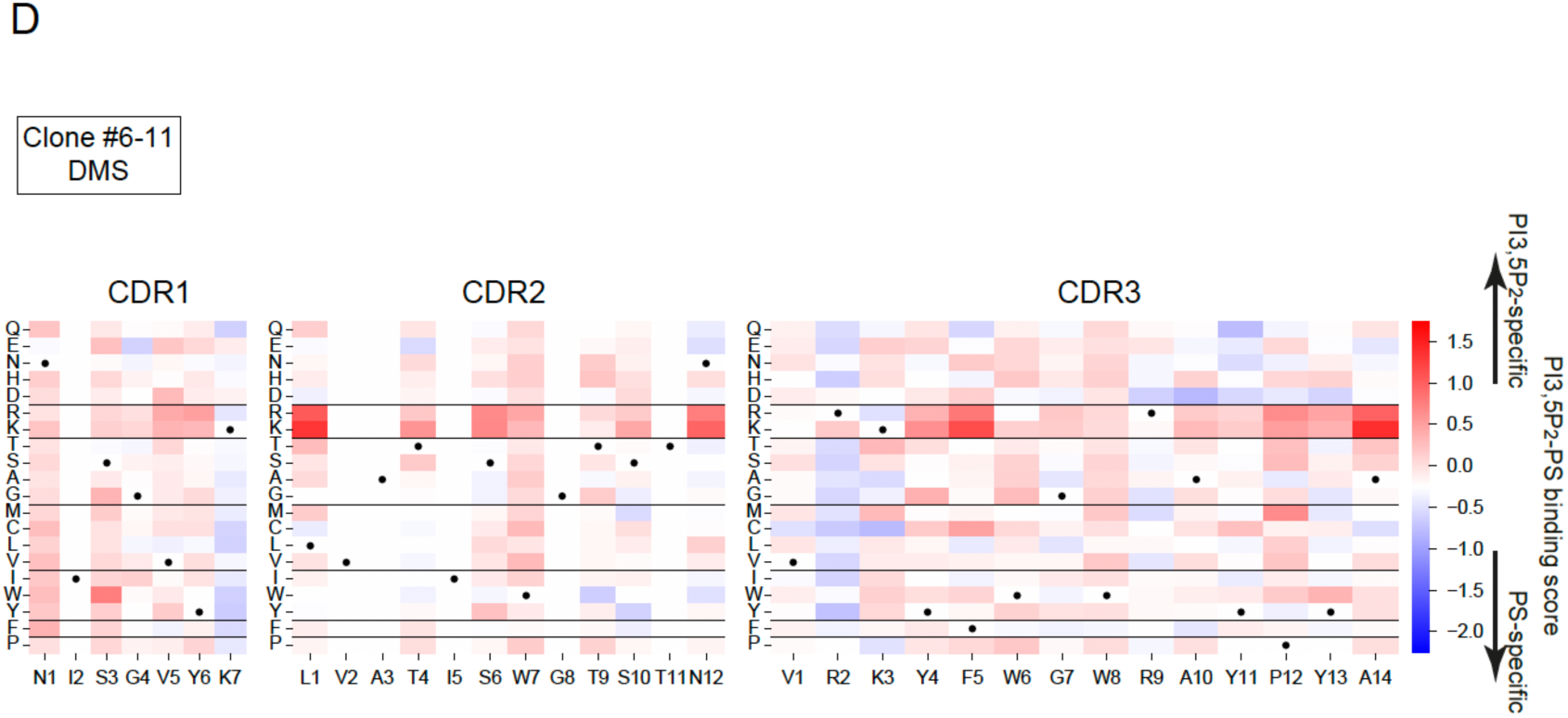
Analysis of isolated PIPs-binding nanobodies. (**A**) The analysis of lipid-binding activities of the isolated nanobody clones by CLiB assay. The BJ5465 protease-deficient strain was used. Data represent mean ± SEM (n = 3). The amino acid sequences of the nanobody clones are listed in Table S2. (**B, C**) The analysis of the lipid-binding activities of the sorted fractions in CLiB assay using the indicated DMS libraries and liposomes. X axis and Y axis indicate the surface expression level and lipid-binding activity of the nanobodies, respectively (B). The histogram of rhodamine signals in the HA-positive cells incubated with the indicated liposomes (C). (**D**) Binding specificity score (PI(3,5)P_2_ [target liposome] binding score - PS [non-target liposome] binding score) distribution of the single point mutants of nanobody clone #6-11. Red and blue indicate clones preferentially binding to PI(3,5)P_2_ and PS, respectively.

**Fig. S5.**
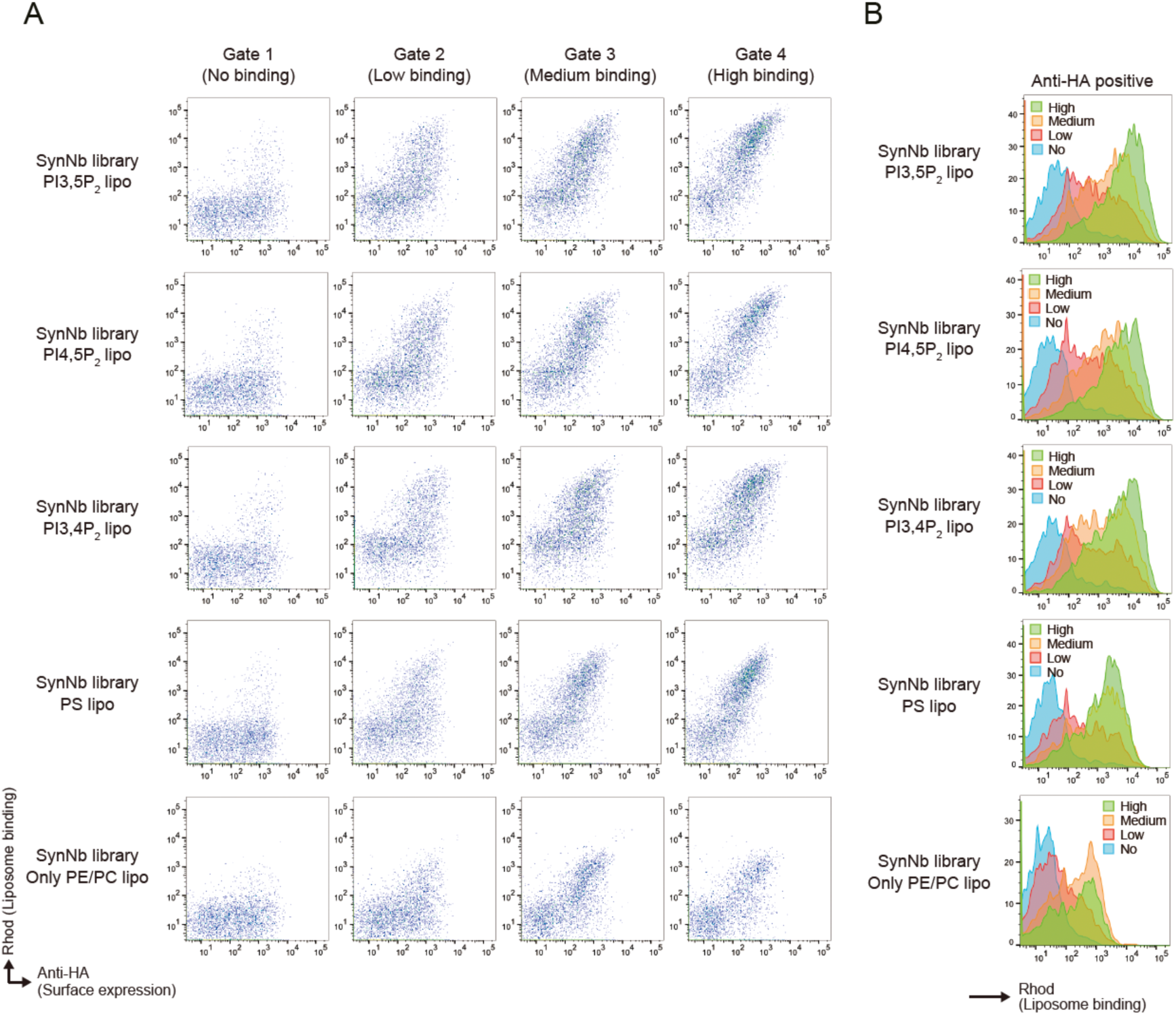
Lipid-binding analysis of each fraction isolated in the DMS analysis of nanobodies. (**A, B**) The analysis of the lipid-binding activities of the sorted fractions in CLiB assay using the indicated DMS libraries and liposomes. X axis and Y axis indicate the surface expression level and lipid-binding activity of the nanobodies, respectively (A). The histogram of rhodamine signals in the HA-positive cells incubated with the indicated liposomes (B).

**Table S1.**
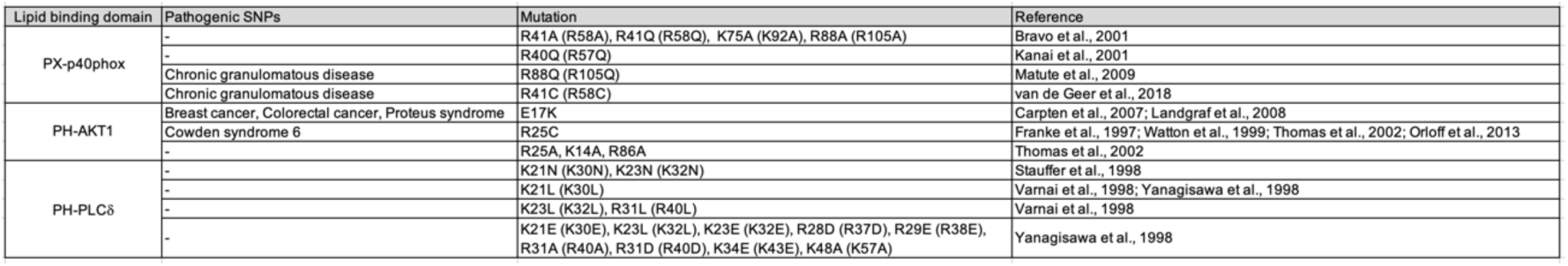
Various mutations in lipid-binding domains reported in the previous studies. The numbers in brackets indicate the position of the original amino acid residues shown in the references.

**Table S2.**
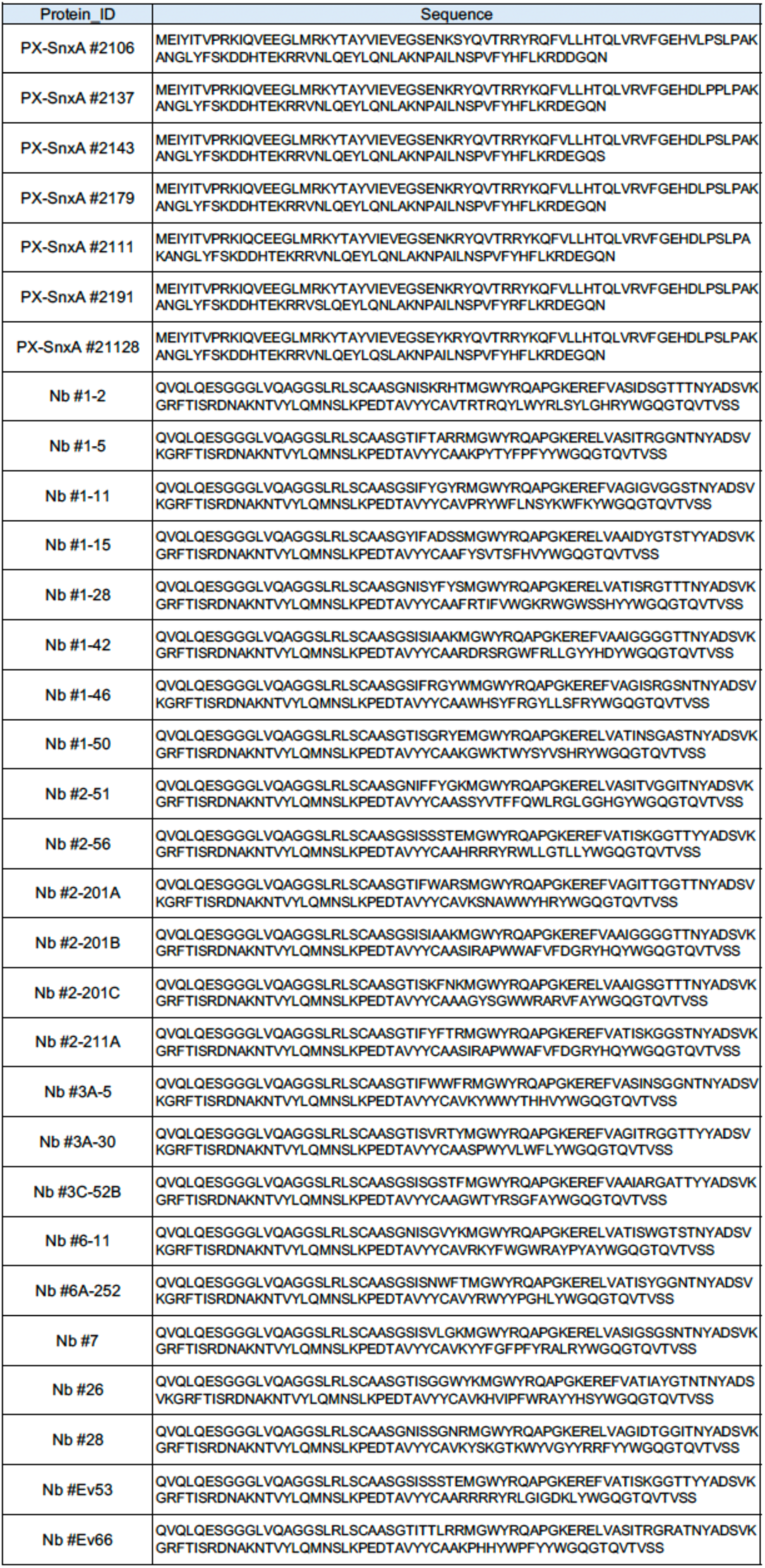
Amino acid sequences of mutated PX-SnxA clones and the isolated nanobody clones.

**Table S3.**
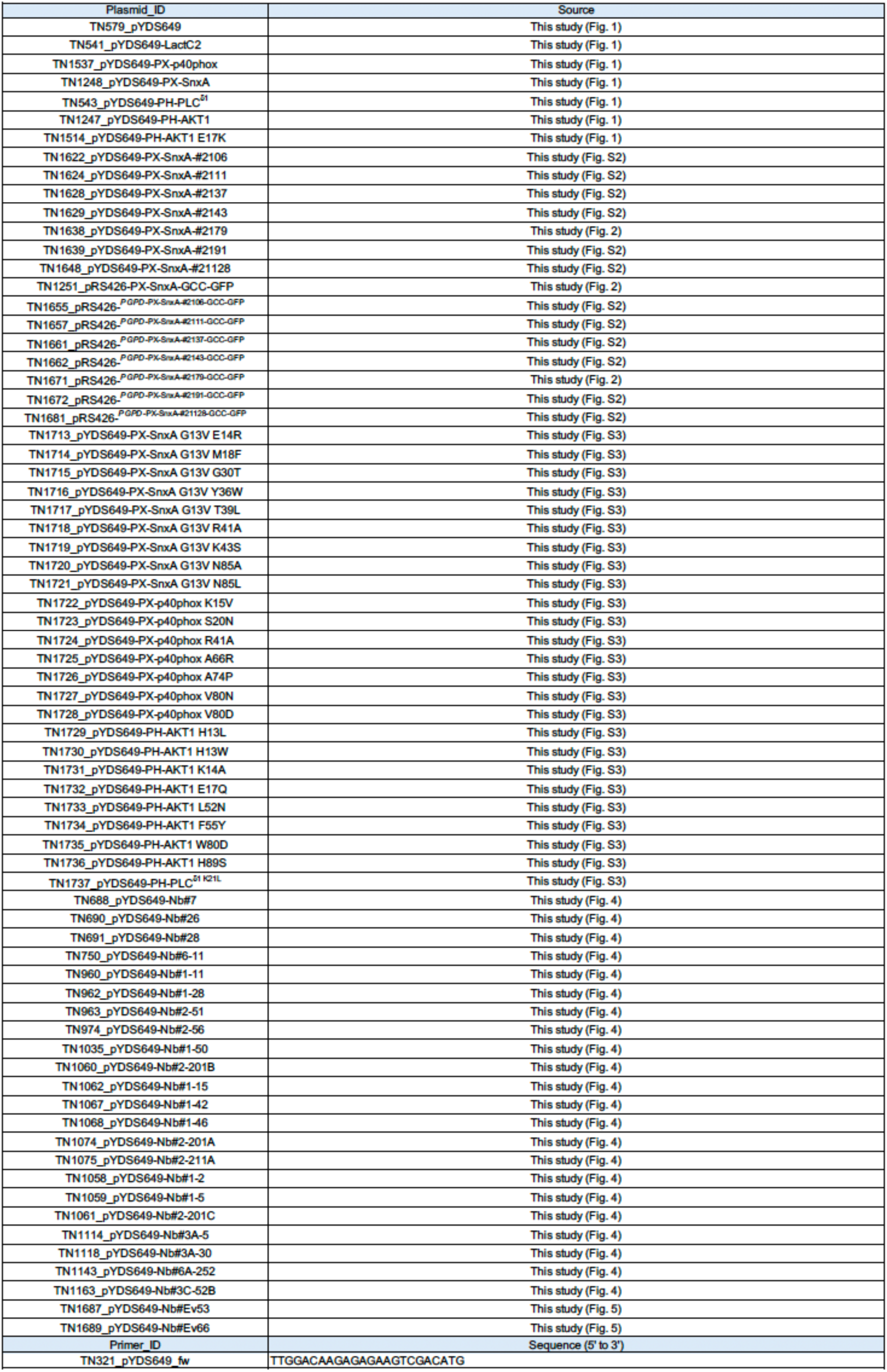
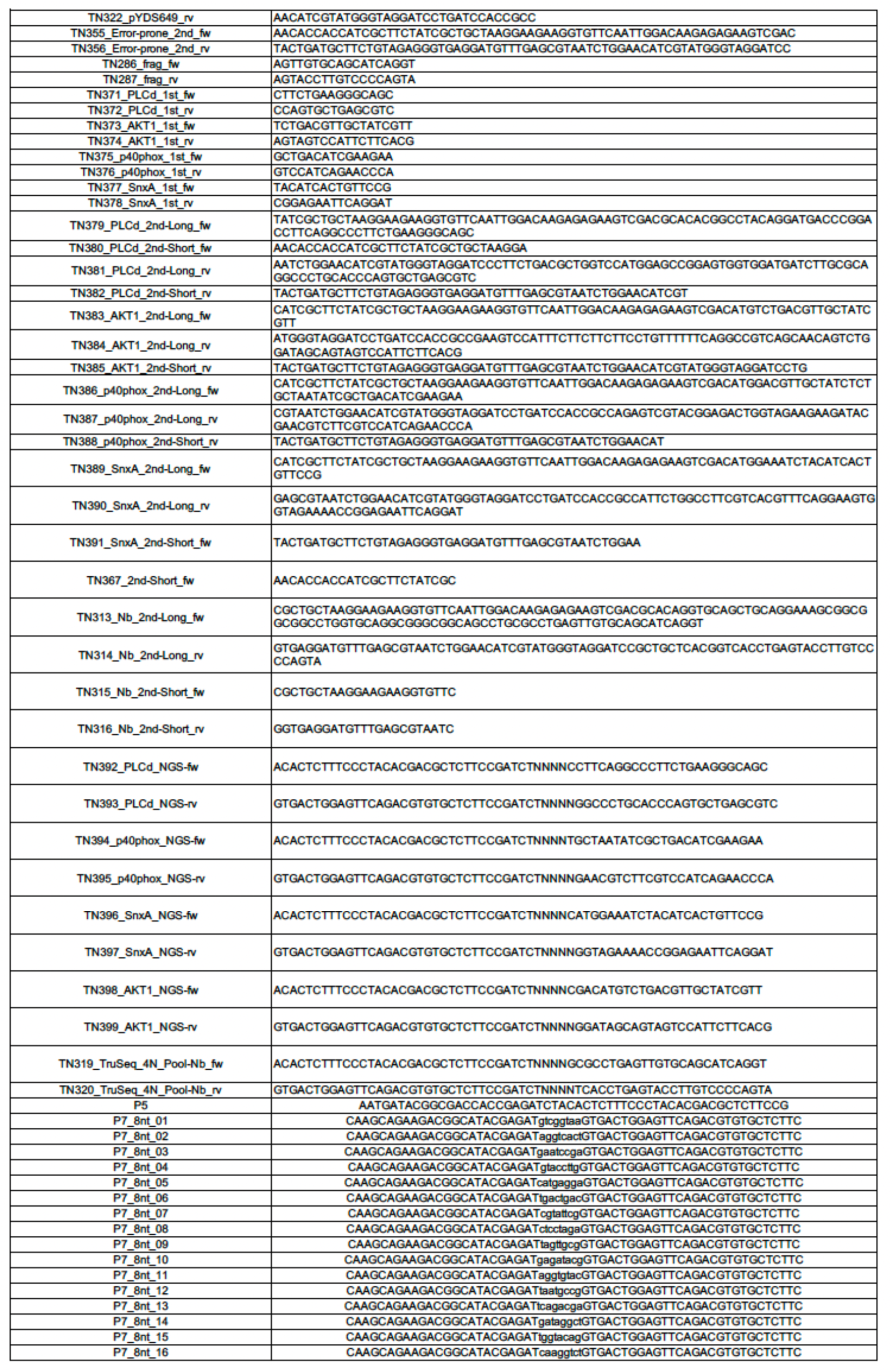
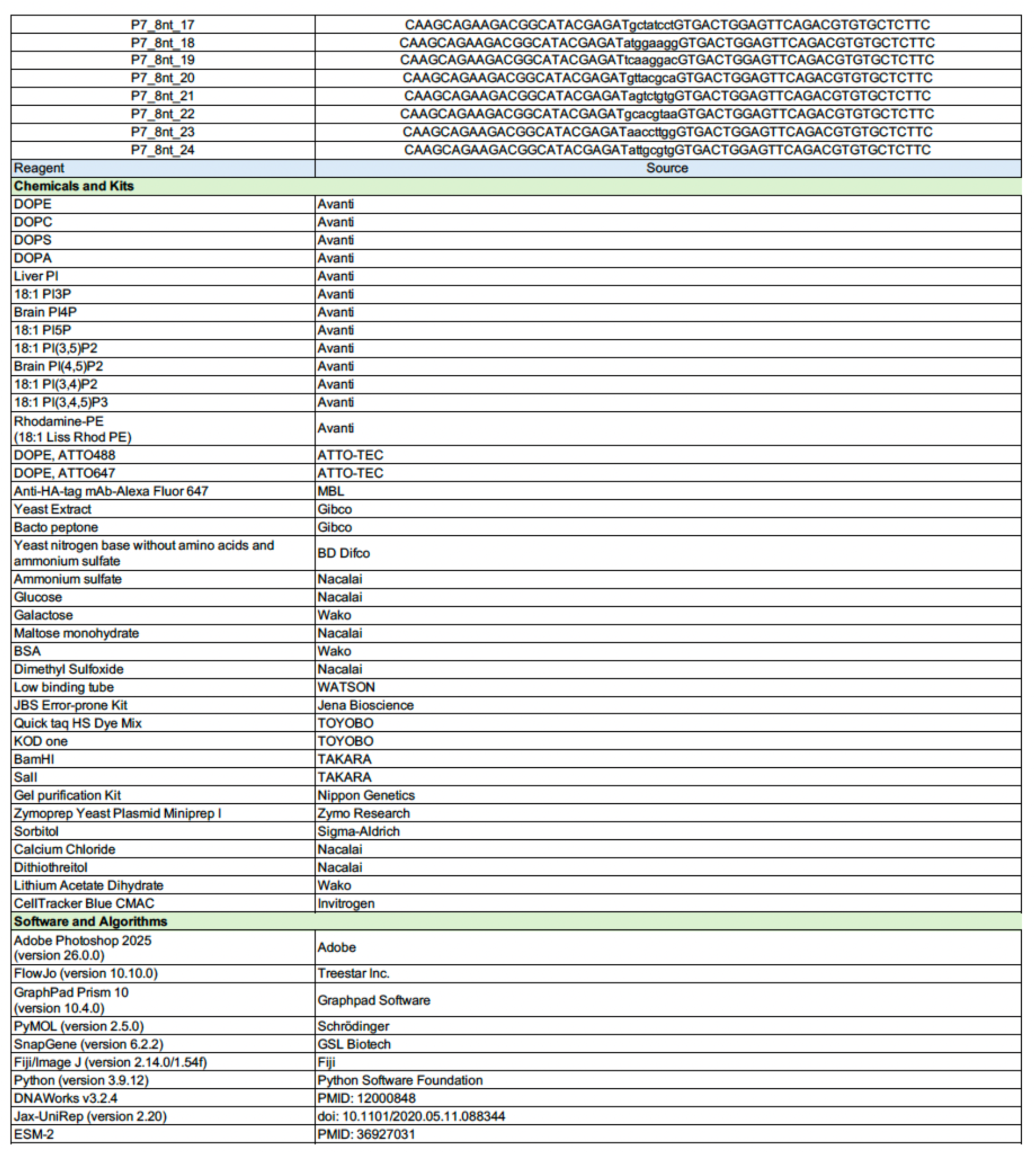
Plasmids, primers and materials used in this study.

**Table S4.** Yeast cell lines used in this study.

| Yeast ID | Yeast line | Source | Description |
| --- | --- | --- | --- |
| YTN61 | SEY6210 [*MATα leu2-3,112 ura3-52 his3Δ200 trp1-Δ901 lys2-801 suc2Δ9*] | PMID: 3062374 | For CLiB assay |
| YTN132 | BJ5465 [*MATα ura352 trp1 leu2Δ1 his3Δ200 pep4::HIS3 prb1Δ1.6 R can1 GAL*] | ATCC | For CLiB assay & HT-CLiB assay |
| YTN161 | 20231211A<br>Mutated PX-SnxA library (BJ5465) | This study (Fig. 2) | For directed evolution<br>2.38 x 10<sup>9</sup> diversity<br>5.9 x 10<sup>9</sup> cells/ml |
| YTN162 | 20240717D<br>PX-SnxA G13V DMS library (BJ5465) | This study (Fig. 3) | For HT-CLiB assay<br>3 x 10<sup>7</sup> diversity<br>1.83 x 10<sup>9</sup> cells/ml |
| YTN163 | 20240717C<br>PX-p40phox DMS library (BJ5465) | This study (Fig. 3) | For HT-CLiB assay<br>1 x 10<sup>7</sup> diversity<br>1.78 x 10<sup>9</sup> cells/ml |
| YTN164 | 20240717B<br>PH-AKT1 DMS library (BJ5465) | This study (Fig. 3) | For HT-CLiB assay<br>6 x 10<sup>7</sup> diversity<br>1.97 x 10<sup>9</sup> cells/ml |
| YTN165 | 20240717A<br>PH-PLCδ1 DMS library (BJ5465) | This study (Fig. S3) | For HT-CLiB assay<br>5 x 10<sup>7</sup> diversity<br>1.84 x 10<sup>9</sup> cells/ml |
| YTN166 | Synthetic Nanobody<br>Yeast Display Library | PMID: 29434346<br>(Kerafast, EF0014-FP) | For nanobody screening<br>2.0 x 10<sup>10</sup> cells/ml |
| YTN167 | 20230630<br>Nanobody DMS library (BJ5465) | This study (Fig. 4) | For HT-CLiB assay<br>4.9 x 10<sup>8</sup> diversity<br>2.0 x 10<sup>9</sup> cells/ml |
| YTN168 | 20240717E<br>In silico designed nanobody library (BJ5465) | This study (Fig. 5) | For HT-CLiB assay<br>4 x 10<sup>7</sup> diversity<br>1.89 x 10<sup>9</sup> cells/ml |

